# The displacement of the σ^70^ finger in initial transcription is highly heterogeneous and promoter-dependent

**DOI:** 10.1101/2023.06.10.544452

**Authors:** Anna Wanga, Abhishek Mazumdera, Achillefs N. Kapanidis

## Abstract

Bacterial sigma factors (σ) contain a highly conserved structural module, the ‘‘σ-finger’’, which forms a loop that protrudes towards the RNA polymerase (RNAP) active-centre in the open complex and has been implicated in pre-organisation of template DNA, abortive initiation of short RNAs, initiation pausing, and promoter escape. Here, we introduce a novel single-molecule FRET (smFRET) assay to monitor σ-finger motions during transcription initiation and promoter escape. We find that the σ-finger is displaced from its position inside the active site cleft before promoter escape, and after synthesis of RNAs with lengths that are highly dependent on the sequence of the promoter used. Real-time smFRET measurements reveal the presence of significant heterogeneity in the timing of finger displacement and show that different σ-finger conformations in single open transcription complexes are associated with substantially different kinetics in transcription initiation and promoter escape, potentially impacting gene regulation in bacteria.

## INTRODUCTION

All cellular core RNA polymerases (RNAP) require at least one protein factor to perform promoter-specific transcription initiation^1–3^. In bacterial transcription, the core RNAP associates with a σ-factor (either the primary-σ [σ^70^] or alternative σ-factors [such as σ^24^, σ^28^, σ^32^, σ^38^, and σ^54^]) to carry out promoter-specific transcription initiation^1^. Nearly all RNAP-σ holoenzymes (except RNAP-σ^54^; see below) contain a structurally similar module called the “σ-finger” (also referred to as σR3-σR4 linker or σR2-σR4 linker), which is positioned inside the active centre cleft of RNAP and interacts with unwound template strand DNA^4–8^. The RNAP-σ^54^ contains a structurally unrelated module, RII.3, which occupies a position analogous to that of the σ-finger module, and has been hypothesised to play a similar functional role^9^. Remarkably, all three domains of life feature σ-finger-like structural elements which protrude into the RNAP active-centre cleft to interact and stabilise the unwound template-strand DNA in conformations that facilitate the binding of initiating nucleotides^4–17^.

Observation of high-resolution structures of bacterial transcription initiation complexes showed that the σ-finger is found along the path of the growing RNA chain (Fig. 1A); it has thus been postulated for a long time that the σ-finger needs to be displaced from the path of the RNA to allow for productive transcription, with the displacement thought to occur either before or during RNA synthesis in initial transcription^4, 18–20^. Consistent with the structural studies, biochemical studies have reported altered transcription profiles for RNAP-σ holoenzymes containing σ-finger mutations or deletions^21, 22^; further, a recent single-molecule study on the kinetics of initial transcription reported significant changes in pausing profiles during initial transcription by an RNAP derivative with a σ-finger deletion^23^.

**Fig. 1.**
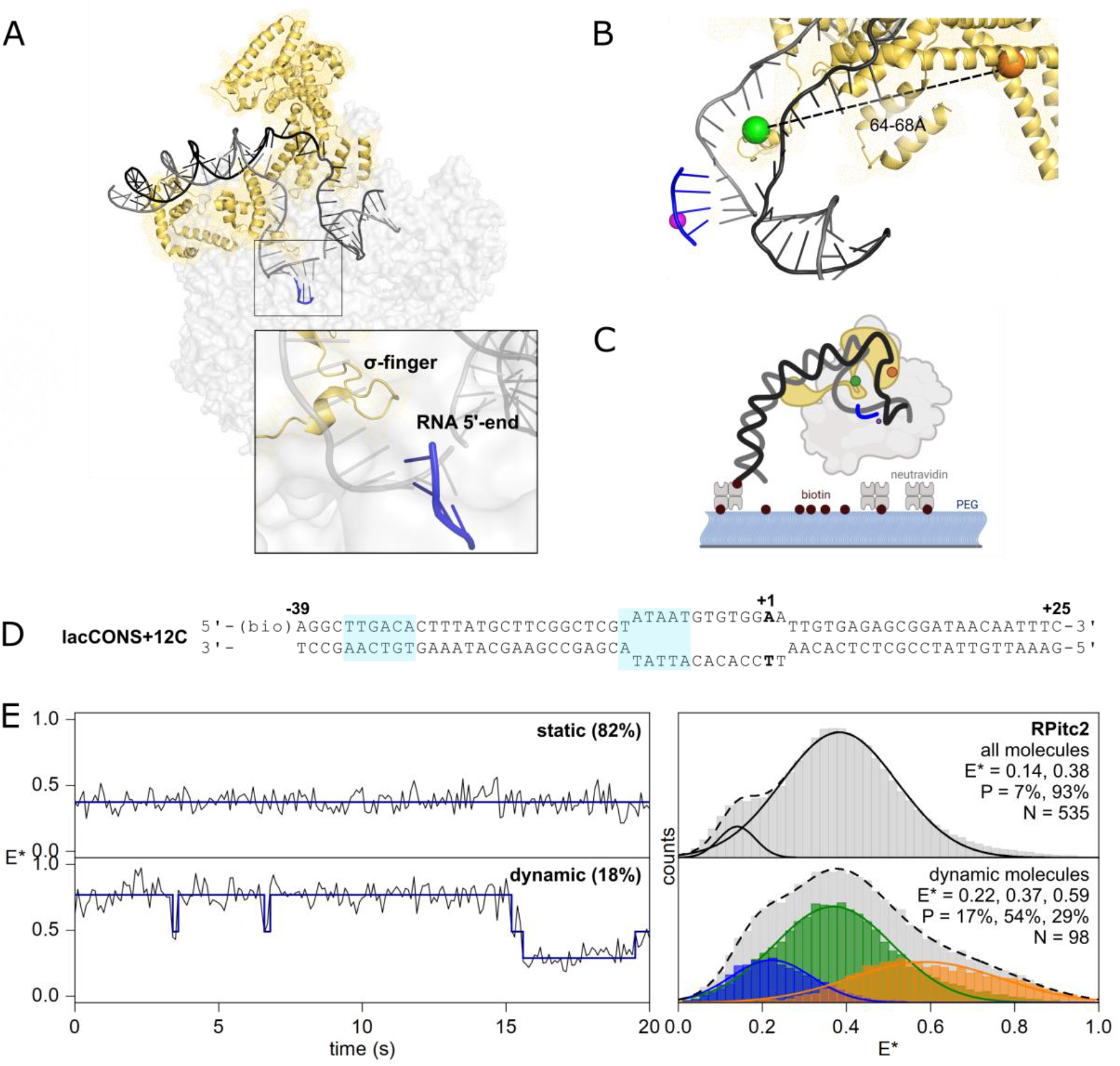
An smFRET assay for detecting σ-finger movements in solution. (A) Conformational state of the pre-displaced σ-finger in *E. coli* RP_itc4_ (PDB 4YLN). Grey, black and blue ribbons show the template DNA strand, non-template DNA strand and transcribed RNA, respectively. Straw ribbons and mesh show σ^70^. (B) smFRET labelling scheme. Orange and green spheres: fluorescent probes at residues 511 and 366 of σ^70^. Purple sphere: catalytic Mg^2+^ ion. (C) Schematic of the assay used for experiments tracking σ-finger movements. Colour scheme as in panel (A) but with the addition of RNAP core shown in light grey. (D) lacCONS DNA fragment used for the assay to track σ-finger movements at RP_itc2_. (E) smFRET data for the σ-finger in RP_itc≤2_ complexes formed using ApA dinucleotide primer showing static (*upper*) and dynamic (*lower*) behaviour. Left, representative traces of static and dynamic behaviour. Right, E* histograms formed as a result of hidden Markov modelling, and Gaussian fitting of sub-populations.

Further insight on σ-finger interactions and conformations during initiation was provided by a recent study that reported several high-resolution structures of *T. thermophilus* RNAP and *M. tuberculosis* RNAP transcription initiation complexes and defined σ-finger interactions with a DNA-RNA hybrid containing different lengths of RNA^18^. The reported structures indicated stabilisation of very short DNA-RNA hybrids, also reported in Brodolin et al.^24^, and stepwise displacement of the tip of the σ-finger from the initial position in the active site cleft as the RNA length increases (>4 nt) and the RNA clashes with the σ-finger. Based on these structures, it has been hypothesized that the σ-finger acts as a “protein spring” by folding back on itself, storing energy which is utilised later during the large-scale disruption of RNAP-promoter and RNAP-σ interactions that occurs in promoter escape. However, the study used nucleic acid scaffolds which did not contain a scrunched DNA^25^ as would be found in actual initial transcribing complexes. As a result, the exact mechanism and kinetics of σ-finger displacement during active transcription in the context of a full promoter remain unresolved.

In this work, we define the σ-finger conformation and dynamics at different stages of transcription initiation -- the open complex (RP_o_), initial transcribing complexes (RP_itc_), and an elongation complex (RD_e_) -- for three well characterised promoters with different properties: a consensus bacterial promoter based on *lac* promoter (lacCONS), a phage λ promoter (pR) and a ribosomal RNA promoter (rrnBP1). Using single-molecule FRET (smFRET) approaches, we identify a new “displaced” σ-finger conformation, define the point of σ-finger displacement for different promoters, identify determinants of σ-finger displacement, and measure the kinetics of σ-finger displacement during active transcription. Our results show surprising diversity between promoters regarding the RNA-length at which σ-finger displacement occurs and reveals the presence of RP_o_ molecules with an order of magnitude difference in σ-finger displacement kinetics during initial transcription.

## RESULTS

### Strategy for detecting σ-finger motions at the single-molecule level

For the detection of σ-finger motions by smFRET during initial transcription, we designed a doubly labelled hexahistidine-tagged RNAP-σ^70^ holoenzyme (DL RNAP-σ^70^), with one fluorescent probe attached to the σ-finger, and another fluorescent probe attached to σR2 (σ^70^ residue 366) -- a position which remains stable throughout initial transcription^25^. Since the σ-finger is a small structural module which interacts with the growing RNA during initial transcription, the position of attachment of the fluorescent probe was carefully chosen to minimise any interference with RNAP function. Accessible volume (AV) modelling of fluorescent probes^26^ attached to positions through 511 to 521 of σ^70^ on high-resolution structures of initial transcribing complexes having a 3-, 4-, or 5-nt RNA revealed that a fluorescent probe placed at position 511 (located at the *base* of the finger, away from the tip of the finger) should not interfere with either σ-finger interactions and RNA interactions (Fig. S1A). The distance between fluorescent probes Cy3B and Alexa647 placed respectively at positions 511 and 366 of σ^70^ was estimated to be in the 61-66 Å range (using PDB structures 4YLN, 7KHB, 7MKD and 7MKE), with corresponding estimated FRET efficiencies (E) in the 0.3-0.5 range (Fig. 1B) – a range well suited for monitoring distance changes of ∼5-20 Å. We thus selected these two positions to generate a DL RNAP-σ^70^ construct (see *Methods*; Fig. S2). The transcriptional activity of DL RNAP-σ^70^ was >80% compared to the activity of unmodified RNAP-σ^70^, indicating that the attached fluorescent probes did not significantly affect RNAP assembly and RNAP function. *In-vitro* transcription assays also showed that the initial transcription profiles obtained using the DL RNAP-σ^70^and a wild-type RNAP-σ^70^ were similar, indicating that the introduction of a fluorescent probe at the base of the finger did not alter the function of the initial transcribing complex (Fig. S3).

### The σ-finger is mobile in the RNAP holoenzyme and in an RNAP-promoter complex

To assess σ-finger conformations in transcription initiation, we incubated DL RNAP-σ^70^ with a biotinylated consensus bacterial core promoter fragment (biotin-*lacCONS;* Table S1, and Fig. 1D) to form heparin-resistant RNAP-promoter open complexes (RP_o_) using established protocols (see *Methods*), followed by immobilisation of individual RP_o_ molecules to glass slides functionalised with neutravidin (Fig. 1C). Next, we added the initiating dinucleotide ApA to the immobilised RP_o_ molecules to form an initial transcribing complex (RP_itc2_) and carried out total internal reflection fluorescence (TIRF) microscopy with alternating laser excitation (ALEX) experiments as described previously^23, 27, 28^. The data were analysed to obtain apparent smFRET efficiencies (E*) from E*-vs-time trajectories of single RP_itc2_ complexes, and to generate an E* histogram that reports on the distribution of σ-finger conformational states.

The E* histogram for RP_itc2_ was multimodal with two subpopulations (Fig. 1E, top right): a major subpopulation with mean E*∼0.38 (93%), and a minor subpopulation with mean E*∼0.14 (7%). Inspection of E*-vs-time trajectories (Fig. 1E, left) revealed a subpopulation of molecules (18%) which showed transitions between different E* values; analysis of E*-vs-time trajectories from these dynamic molecules using Hidden Markov Modelling^29, 30^ identified three subpopulations with mean E* of ∼0.22 (17%), ∼0.37 (54%), and ∼0.59 (29%) and dwell times in the range of ∼0.6-1.0 s (Fig. 1E, bottom right).

Distance estimates corresponding to subpopulations with mean E* values of ∼0.37 (corresponding to distances of ∼66Å) were in excellent agreement to those predicted from AV calculations on high-resolution structures of *E. coli* RNAP and RNAP-promoter complexes (65.9±9.1Å, 4YLN; 61.4±8.2Å, 7KHB; 64.7±8.9Å, 7MKD; and 62.7±6.1Å, 7MKE). We thus assigned subpopulations with mean E* values of ∼0.37 to “in-cleft” σ-finger conformations positioned to interact with a single-stranded template DNA loaded in the RNAP main channel as visualised in the high-resolution structures. On the other hand, subpopulations with mean E* values of ∼0.22 represent structural states where the σ-finger is positioned away from σR2, whereas subpopulations with mean E*of ∼0.59 represent structural states where the σ-finger is positioned closer to σR2.

In a next set of experiments, we measured apparent smFRET efficiencies (E*) of DL RNAP-σ^70^ molecules immobilised via a hexahistidine tag on a glass slide functionalized with anti-hexahistidine tag antibody (Fig. S4A). Use of ALEX allows selection of DL RNAP-σ^70^ molecules with a single donor-acceptor probe pair and measurement of corresponding E* time trajectories. As for RP_itc2_, the overall E* histogram for DL RNAP-σ^70^ revealed a multimodal distribution, with populations centred around mean E* values of ∼0.15 (10%), ∼0.37 (72%) and ∼0.69 (18%) (Fig. S4B). A visual inspection of the E*-vs-time trajectories revealed a class of molecules (28%) which exhibited transitions between different FRET states and were classified as dynamic molecules, within which we identified three subpopulations having mean E* values of ∼0.22 (28%), ∼0.45 (54%), and ∼0.70 (18%), all with dwell times in these conformational states on time scales of ∼0.5-0.8 s (Fig. S4C). Overall, our results were similar for DL RNAP-σ^70^ and RP_itc2_, revealing multiple σ-finger subpopulations and significant σ-finger conformational dynamics in these complexes.

### σ-finger displacement during initial transcription: consensus bacterial promoter (lacCONS)

We then assessed σ-finger conformations in primer-dependent initial transcription for synthesis of RNAs up to 11-nt in length. Initial transcribing complexes (RP_itc_) were formed by adding subsets of nucleotides to RP_itc2_ complexes prepared using lacCONS constructs (Table S1). Use of subsets directed transcription up to a desired maximum RNA length when CTP is withheld; further, use of ApA dinucleotide as a primer led to synthesis of RNAs with a 5’-OH group.

Analysis of E*-vs-time trajectories for experiments for RP_itc≤4_ resulted in E* histograms similar to those for RP_itc2,_ i.e., showing two subpopulations with E*∼0.41 (93%) and E*∼0.14 (7%; Fig. 2A); however, unlike the results for RP_itc2_, a much smaller fraction of molecules showed dynamics (9%; Fig. 2D, green line).

**Fig. 2.**
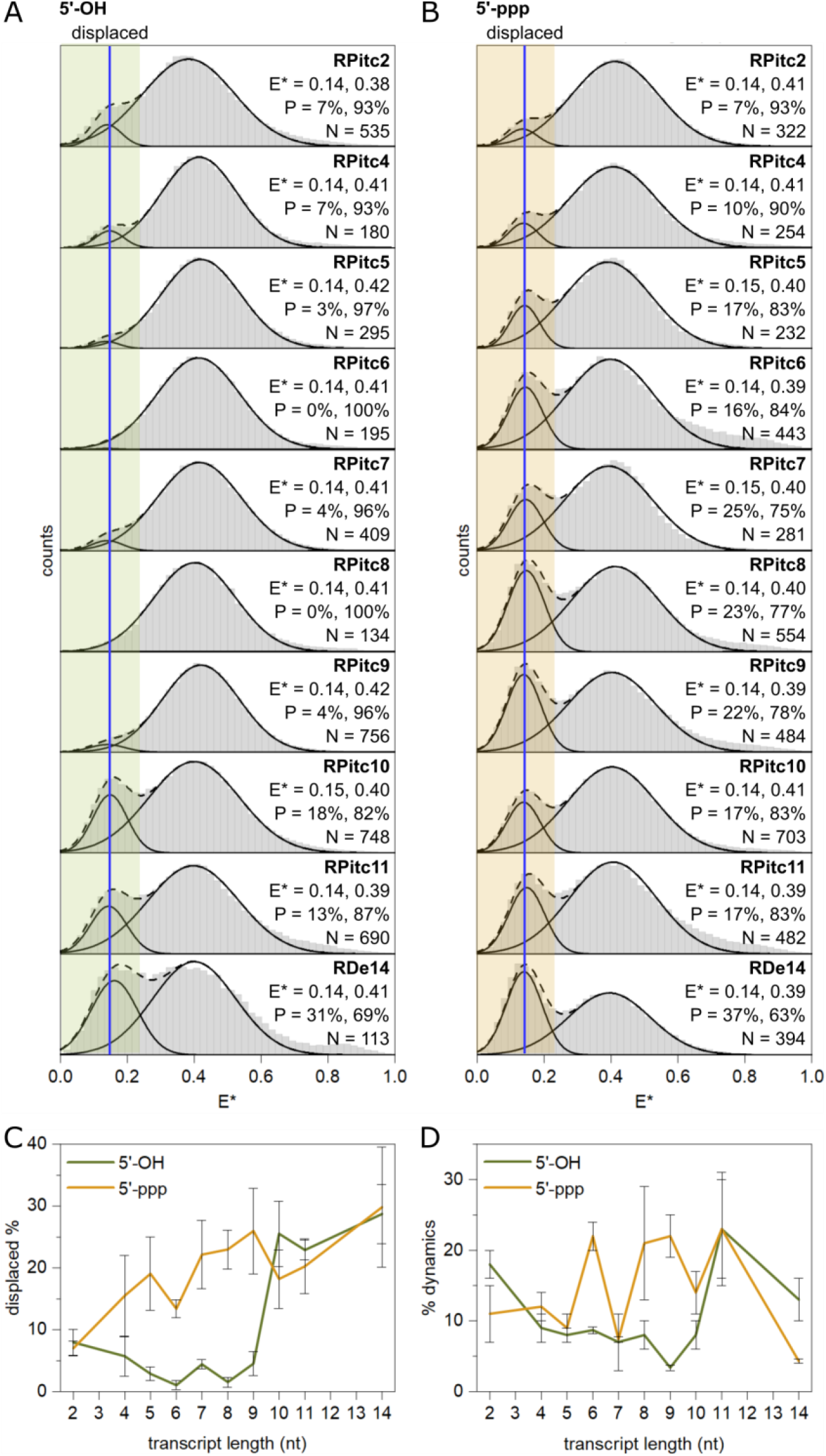
σ-finger displacement is affected by the 5’-end of RNA. (A) E* histograms of the σ-finger at stages of transcription involving the lacCONS promoter and RNA with a 5’-hydroxyl end (2, 4, 5, 6, 7, 8, 9, 10 and 11 nucleotides in length). Estimations of mean E* values and occupation probabilities are shown for the displaced (blue line, green highlighting) and pre-displaced populations. (B) E* histograms of the σ-finger at stages of transcription involving the lacCONS promoter and RNA with a 5’-triphosphate end (2, 4, 5, 6, 7, 8, 9, 10 and 11 nucleotides in length). Estimates of mean E* values and occupation probabilities are shown for the displaced (blue line, orange highlighting) and pre-displaced populations. (C) Occupation probabilities of the displaced σ-finger conformation with transcript length for RNA with a 5’-hydroxyl (green line) and a 5’-triphosphate (orange line) end. Error bars are calculated from the standard deviation in the mean between different subsets and repeats of the experiment. (D) Percentage of dynamic molecules with transcript length for RNA with a 5’-hydroxyl (green line) and a 5’-triphosphate (orange line) end. Error bars are calculated from the standard deviation in the mean between different subsets and repeats of the datasets.

For complexes making RNAs of up to 9-nt in length (complexes ranging from RP_itc≤5_ to RP_itc≤9_), the E* histograms showed almost exclusively the presence of the subpopulation corresponding to a “in-cleft” σ-finger conformation (96%-100%; Fig. 2A). Significantly, the E* time trajectories showed few dynamics (<10% of dynamic molecules; Fig. 2D), indicating formation of a stabilised conformation for the σ-finger module during this stage of initial transcription. The lack of significant structural heterogeneity and dynamics for our construct (which probes the motions of the *base* of the σ-finger; Fig. S1B) indicates that it forms a stable structural state for the base of the finger when a 5-mer RNA is synthesised, and stays in similarly stable states for RNA extensions up to a 9-mer.

In contrast, further RNA extension beyond 9-nt led to increased abundance for low-FRET species (E*∼0.15 (19%) for RP_itc≤10_ and E*∼0.14 (18%) for RP_itc≤11_; Fig. 2A), indicating that, for a significant fraction of molecules, synthesis of a 10-mer RNA results in σ-finger displacement from an “in-cleft” conformation to a conformation positioned away from σR2, which we assign to a “displaced” σ-finger conformation.

Notably, analysis of E* time trajectories for RP_itc≤11_ revealed a much higher fraction of dynamic molecules (21%, Fig. 2D), with transitions between three subpopulations centred around E*∼0.23, 0.42 and 0.64, occurring on timescales of ∼1.0-1.4 s (Fig. S5). The increased mobility of the σ-finger for a large fraction of molecules during late stages of initial transcription for lacCONS (11-nt RNA) as compared to earlier stages (4-10 nt RNA) may originate either from a loss of contact with the scrunched DNA-RNA hybrid (resulting in motions similar to the ones observed in the early stages of initial transcription, i.e., RP_itc2_), or due to motions of the entire σ-finger DNA-RNA hybrid associated with a “unscrunching-scrunching” of the transcription bubble, as reported in previous single-molecule studies^31^. If the resulting dynamics were indeed similar to that observed in the early stages of initial transcription, then this result would suggest that, at the stage of finger displacement, the σ-finger breaks its contact with the DNA-RNA hybrid and becomes mobile, exploring several conformational states. However, a dwell-time analysis for subpopulations with a mean E*∼0.23 revealed a peaked distribution for dwell times corresponding to different conformational states populated in RP_itc≤11;_ this distribution shape is in sharp contrast to exponential distributions observed previously for RP_itc2_ (Fig. S6). A peaked dwell-time distribution suggests a conformational change involving more than one step, indicating that the σ-finger motions in these late stages of initial transcription correlate with the previously observed dynamics of the transcription-bubble^31^.

Further extension to a 14-mer RNA (RD_e≤14_) resulted in a E* histogram with a subpopulation around ∼0.14 (31%) and a reduction in the fraction of dynamic trajectories (to 13%; Fig. 2D). This reduction indicates progression out of the late stages on initial transcription and promoter escape in order to form an early elongation complex that still retains σ^70^ (Ref.32).

To determine the status of the transcription bubble for this promoter during initial transcription, we performed similar experiments using a previously reported construct with donor and acceptor probes at positions −15 of the non-template and +20 of the template DNA^23^ (lacCONS −15/+20; Table S1) and an unlabelled hexahistidine tagged RNAP-σ^70^ holoenzyme. Previous reports for initial transcribing complexes on this construct established that subpopulations with a mean E*∼0.45 represented a scrunched transcription bubble^23^. Our experiments monitoring E*-vs-time trajectories for complexes with 11-mer RNA revealed a large peak around E*∼0.45, indicative of a scrunched bubble which disappeared upon extension to a 14-mer RNA (Fig. S7). The point of escape for the lacCONS promoter therefore lies between the synthesis of an 11-mer and a 14-mer RNA. We note a significant fraction of the complexes remain at E*∼0.45, suggesting that these species include subpopulations of complexes which are either transcriptionally inactive, or engaged in abortive initiation and unable to escape to elongation. Taken together, our data clearly show that, for the lacCONS promoter, the base of the σ-finger is displaced during initial transcription, and before promoter escape.

### Point of σ-finger displacement depends on the functional group at the 5’-end of RNA

We then assessed σ-finger conformations in primer-*independen*t transcription, which results in synthesis of RNAs with a 5’-triphosphate (5’-ppp), a chemical moiety with greater steric bulk and additional negative charges compared to a 5’-OH, and thus with potential for different interactions with the negatively charged σ-finger. We performed experiments similar to those in the previous section but in the absence of a dinucleotide primer, and measured σ-finger conformational dynamics for complexes capable of synthesizing RNAs from up to 2-nt in length (ppp-RP_itc≤2_), to up to 11-nt in length (ppp-RP_itc≤11_). Analysis of E* distributions for ppp-RP_itc≤2_ and ppp-RP_itc≤4_ showed results similar to those obtained for prime-dependent transcription initiation (Fig. 2B). However, further extension to a 5-mer RNA resulted in a E* histogram with an increased fraction of the subpopulation with E*∼0.14 (14%; cf. with 3% for a 5-mer with a 5’-OH end). Similar results were obtained for complexes capable of synthesizing from up to a 6-mer to up to an 11-mer RNA, with occupancies in the E*∼0.14 state being within the ∼18-25% range (Fig. 2B; Fig. 2C, orange line); additionally, a significant fraction (∼14-23%) of E* time trajectories for all but one of these complexes show dynamic behaviour (Fig. 2D, orange line), analogous to results obtained for RP_itc≤11_ in the primer-dependent transcription initiation. Our results indicate that, in primer-independent initiation, the unfavourable interactions between the bulky, negatively charged triphosphate group and the σ-finger (which has a negative net charge) results in displacement at a much shorter RNA length compared to primer-dependent initiation.

### Kinetics of σ-finger displacement during initial transcription

To observe the kinetics of σ-finger displacement during initial transcription, we performed real-time RNA synthesis up to the point of promoter escape during the recording of our single-molecule movies. Specifically, we pre-formed RP_o_ with DL RNAP-σ^70^ and lacCONS promoter DNA fragments at 37°C, immobilised RP_o_ molecules (as in Fig. 1C) and formed RP _itc≤2_ on glass, started recording of the movie, and added a subset of NTPs directing transcription up to the formation of a stable elongation complex (for lacCONS, this corresponds to synthesis of a 14-nt RNA; Fig. 3A). To compare the kinetics of σ-finger displacement during primer-dependent transcription initiation with those of primer-independent transcription initiation, RP_itc2_ was formed with either an ApA dinucleotide primer, or a pppApA dinucleotide primer, respectively. Movies were analysed to extract E* time trajectories and were fit using HMM to extract the time between nucleotide addition and σ-finger displacement (see *Methods*), a time which we will referred to as displacement time t_d_.

**Fig. 3.**
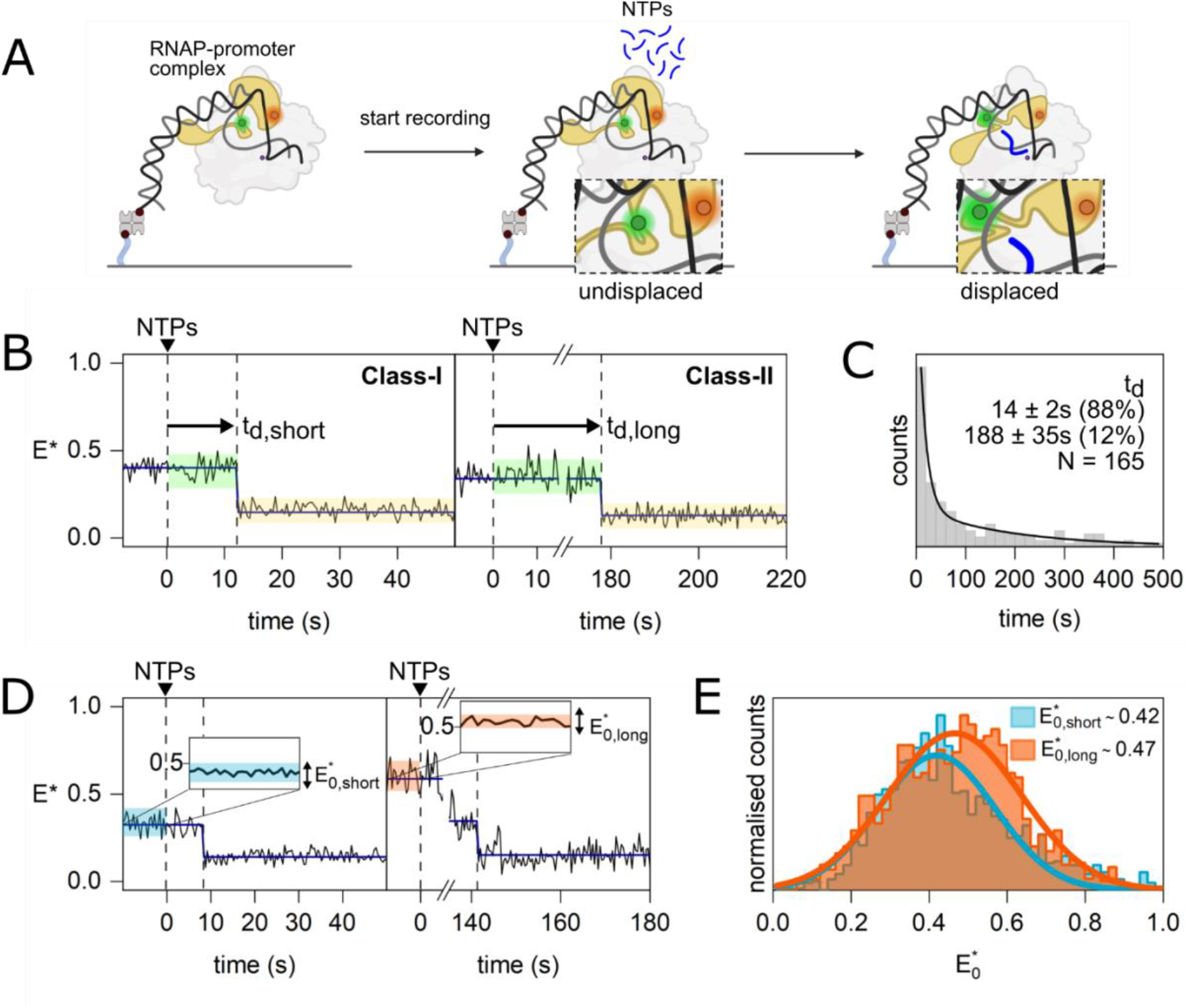
Real-time experiments monitoring the σ-finger conformation in initial transcription complexes containing the lacCONS promoter and RNA with a 5’-OH end. (A) Schematic of the real-time assay used to track σ-finger movements. (B) Example E*-time trajectories of Class-I and Class-II molecules showing σ-finger displacement. The conformation between NTP addition and displacement is highlighted in green, and the conformation after displacement is highlighted in yellow. (C) σ-finger displacement time histogram fitted with a double-exponential decay (black line) showing that displacement occurs on heterogeneous timescales. (D) Example E*-time trajectories of Class-I and Class-II molecules. The time before NTP addition is highlighted for blue and red for Class-I and Class-II molecules respectively. (E) Histograms showing the σ-finger conformation before NTP addition of Class-I (E*_0,short_; blue) and Class-II (E*_0,long_; red) molecules.

A visual examination of the E* time trajectories (Fig. S8) obtained in presence of ApA revealed several classes of molecules. Class-I molecules showed a step decrease in E* to a stable state around E*∼0.15, which corresponds to the “displaced” σ-finger; this transition occurred on a short timescale (Fig. 3B, left). In contrast, Class-II molecules transitioned to a stable “displaced” σ-finger conformation on a long timescale (Fig. 3B, right). Class-I and Class-II molecules made up 25% of all molecules. Class-III molecules (44% of all molecules; Fig. S8B) showed no transition to an E*-state characteristic of the “displaced” σ-finger but exhibited transitions between other E* states. An inspection of E*-vs-time trajectories for molecules in RD_e≤14_ revealed very few dynamic molecules (13%; Fig. 2D), indicating that, for some of the dynamic molecules in Class-III, the σ-finger is displaced at later time points and is missed due to premature termination of the time-trajectory. Finally, Class-IV molecules (31% of all molecules, Fig. S8C) showed no transitions and likely consist of either inactive complexes or complexes that remain at early stages of initial transcription.

To estimate the displacement time t_d_, we combined molecules from Class-I and Class-II, binned them into a dwell-time histogram, and fitted the distribution with a biexponential decay (Fig. 3C). Results showed two types of displacement events, a fast one, t_d,short_ ∼14 s (accounting for ∼90% of events) and a slow one, t_d,long_ ∼190 s (accounting for ∼10% of events).

Similar experiments with pppApA revealed E*-vs-time trajectories that contained all four Classes of molecules seen in the absence of the 5’-triphospate group on the RNA, with abundances of 41% for Class-I and Class-II, 27% for Class-III and 32% for Class-IV (Fig. S9). Analysis of displacement times from Class-I and Class-II also showed some heterogeneity in displacement times, with t_d,short_ ∼1.4 s (∼97% of the events) and t_d,long_ ∼70 s (∼3% of the events; Fig. S10A). Strikingly, the displacement events for Classes I and II for transcription with an RNA having a 5’-triphosphate occurred at much faster timescales compared to transcription with an RNA having a 5’-OH. Together with the observation that the σ-finger is displaced at much shorter RNA lengths for primer-independent initiation (5-nt RNA) compared to a primer-dependent initiation (10-nt RNA), this result points to a much lower energy barrier for σ-finger displacement when a triphosphate is present at the 5’-end of the growing RNA, likely resulting from unfavourable interactions between the largely negatively charged σ-finger module (due to the presence of a stretch of four acidic residues in the σ-finger: D513, D514, E515, and D516) and the bulky, negatively charged triphosphate group.

We then explored the intriguing observation of molecules with very different displacement kinetics. Since the real-time reaction starts with the addition of NTPs to immobilised RP _itc≤2_ complexes, we hypothesized that the different kinetics may be associated with conformationally different RP _itc≤2_ molecules, and especially, with different conformations of the σ-finger, which plays key roles in transcription initiation and promoter escape. To test this hypothesis, we analysed the σ-finger conformations *before* NTP addition for two types of molecules: those showing fast displacement kinetics (“fast-displacers”, with t_d_ < 31 s for primer-dependent initiation and t_d_ < 4 s for primer-independent initiation) and those showing slow displacement kinetics (“slow-displacers”, with t_d_ > 31 s for primer-dependent initiation and t_d_ > 4 s for primer-independent initiation). This analysis revealed distinctly different E* histograms for the two types of species on lacCONS: molecules with a wide E* distribution centred around ∼0.42 for fast displacers, and molecules with a wide E* distribution centred around ∼0.47 for slow displacers (Fig. 3D-E). Similar results on lacCONS were obtained for both primer-dependent (5’-OH RNA) and primer-independent transcription initiation (5’-ppp RNA) (Fig. 3E, S10B).

### σ-finger displacement during initial transcription: a phage λ promoter (pR)

We then examined the profile of σ-finger displacement in different promoters, since it is well known that initial transcription and promoter escape are heavily dependent on the promoter and initially-transcribed sequences^32, 33^. We performed analogous experiments with two widely studied and naturally occurring bacterial promoters, the λ pR and the rrnBP1 (Fig. 4A-B), which exhibit significantly different behaviour during initial transcription. The pR promoter is characterised by formation of a very stable RP_o_^34^ (lifetime >10^4^ s), a 13-bp unwound bubble and a transcription start site seven bases downstream of the −10 element. The pR promoter directs accumulation of short RNA products (abortive initiation) during initial transcription and escape from the promoter at an RNA length of 11-nt^34, 35^. Structural studies of RP_o_ formed at the pR have shown that the σ-finger module protrudes into the active site cleft to contact the single-stranded template DNA and is positioned to clash with a growing RNA chain of 4-6 nt in length^19^ (Fig. S11).

**Fig. 4.**
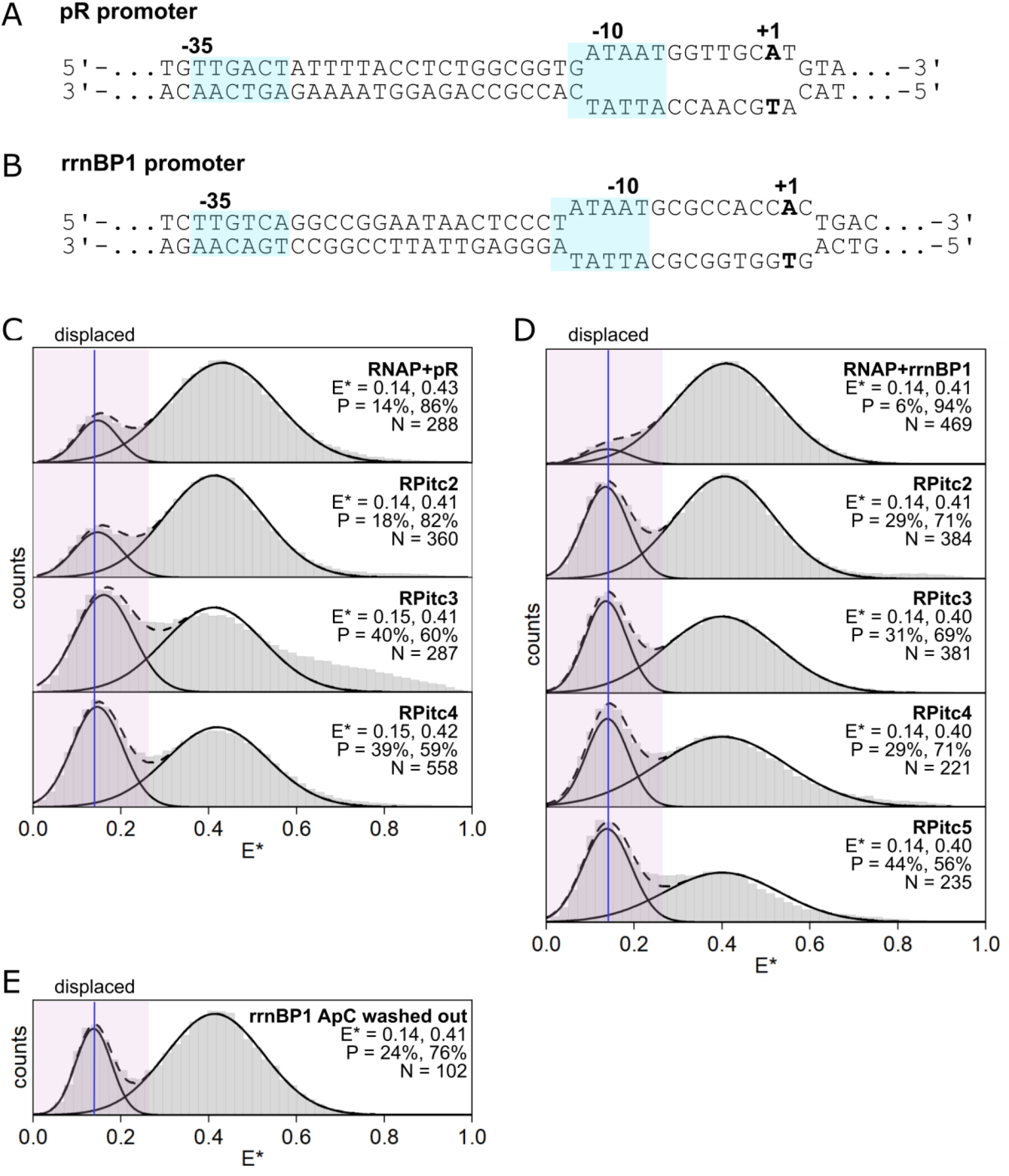
σ-finger displacement depends strongly on the promoter sequence. (A) pR DNA fragment used for the assay to track σ-finger movements. (B) rrnBP1 DNA fragment used for the assay to track σ-finger movements. (C) E* histograms of the σ-finger at stages of transcription involving the pR promoter and RNA with a 5’-hydroxyl end (0, 2 3 4 and 5 nt in length). Estimates of mean E* values and occupation probabilities are shown for the displaced (blue line, purple highlighting) and pre-displaced populations. (D) E* histograms of the σ-finger at stages of transcription involving the rrnBP1 promoter and RNA with a 5’-hydroxyl end (0, 2, 3 and 4 nt in length). Estimations of mean E* values and occupation probabilities are shown for the displaced (blue line, purple highlighting) and pre-displaced populations. (E) E* histogram of the σ-finger conformation involving the rrnBP1 promoter after incubation and removal of ApC.

Experiments performed on an RP_o_ formed on a pR promoter fragment showed a major subpopulation with mean E*∼0.43 (86%) representative of an in-cleft σ-finger, and a minor conformation with mean E*∼0.14 (14%) representative of a displaced σ-finger (Fig. 4C); these results were similar to those obtained on an RP_o_ formed on the lacCONS promoter (Fig. S12). Measurements performed with initial transcribing complexes containing varying lengths of RNA first revealed that the abundance of the subpopulation corresponding to the displaced σ-finger was similar for RP_itc≤2_ (18%). Surprisingly, however, the displaced σ-finger conformation significantly increased for RP_itc≤3_ (40%) and RP_itc≤4_ (39%) (Fig. 4C), indicating that displacement occurs after synthesis of a very short RNA (just a 3-mer), and well before the RNA reaches sufficient length to clash with the σ-finger for the pR promoter (∼5 nt). Further, since it is known that escape for pR promoter occurs during RNA extension from a 10- to 11-mer^34^, our results show that, for the pR promoter, the σ-finger is displaced from the RNAP active site cleft well before promoter escape.

Next, we performed real-time experiments with the pR promoter (Fig. 5A). Our results revealed the presence of two sub-populations of molecules exhibiting short (∼17 s, ∼88% of the events) and long (∼185 s, ∼12% of the events) displacement times (Fig. 5B), essentially identical to the results obtained previously for the lacCONS promoter. However, unlike in lacCONS, an analysis of the E* values for the fast and slow displacers before NTP addition revealed similar E* values (∼0.46 and ∼0.48, respectively; Fig. 5C).

**Fig. 5.**
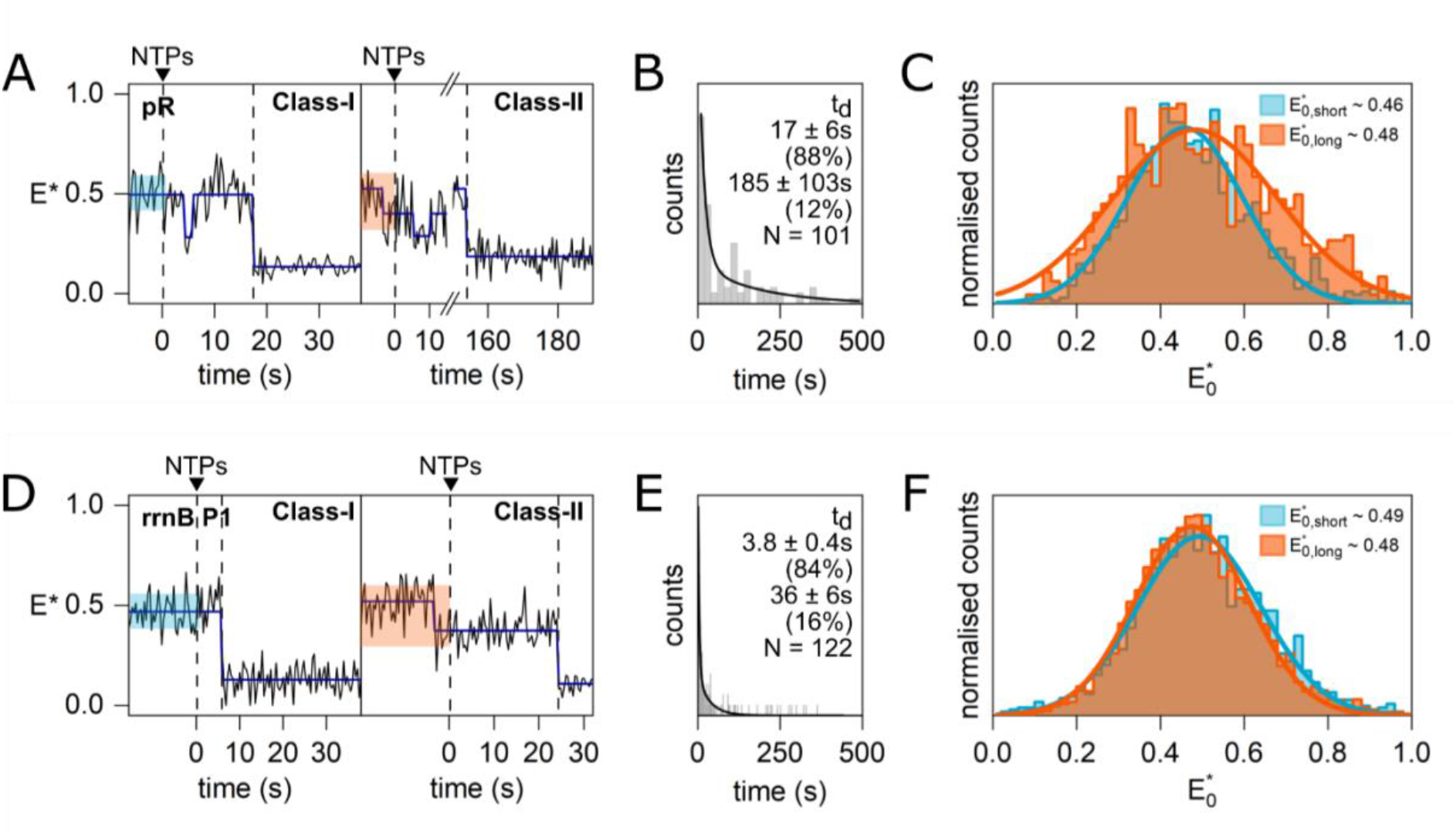
Real-time experiments monitoring the σ-finger conformation in initial transcription complexes containing the pR (A-C) and rrnBP1 (D-F) promoter and RNA with a 5’-OH end. (A, D) Example E*-time trajectories of Class-I and Class-II molecules showing σ-finger displacement. The conformation before NTP addition and displacement is highlighted blue and red for Class-I and Class-II molecules, respectively. (B, E) σ-finger displacement time histogram fitted with a double-exponential decay (black line) showing that displacement occurs on heterogeneous timescales. (C, F) Histograms showing the σ-finger conformation before NTP addition of Class-I (E*_0,short_; blue) and Class-II (E*_0,long_; red) molecules.

**Fig. 6.**
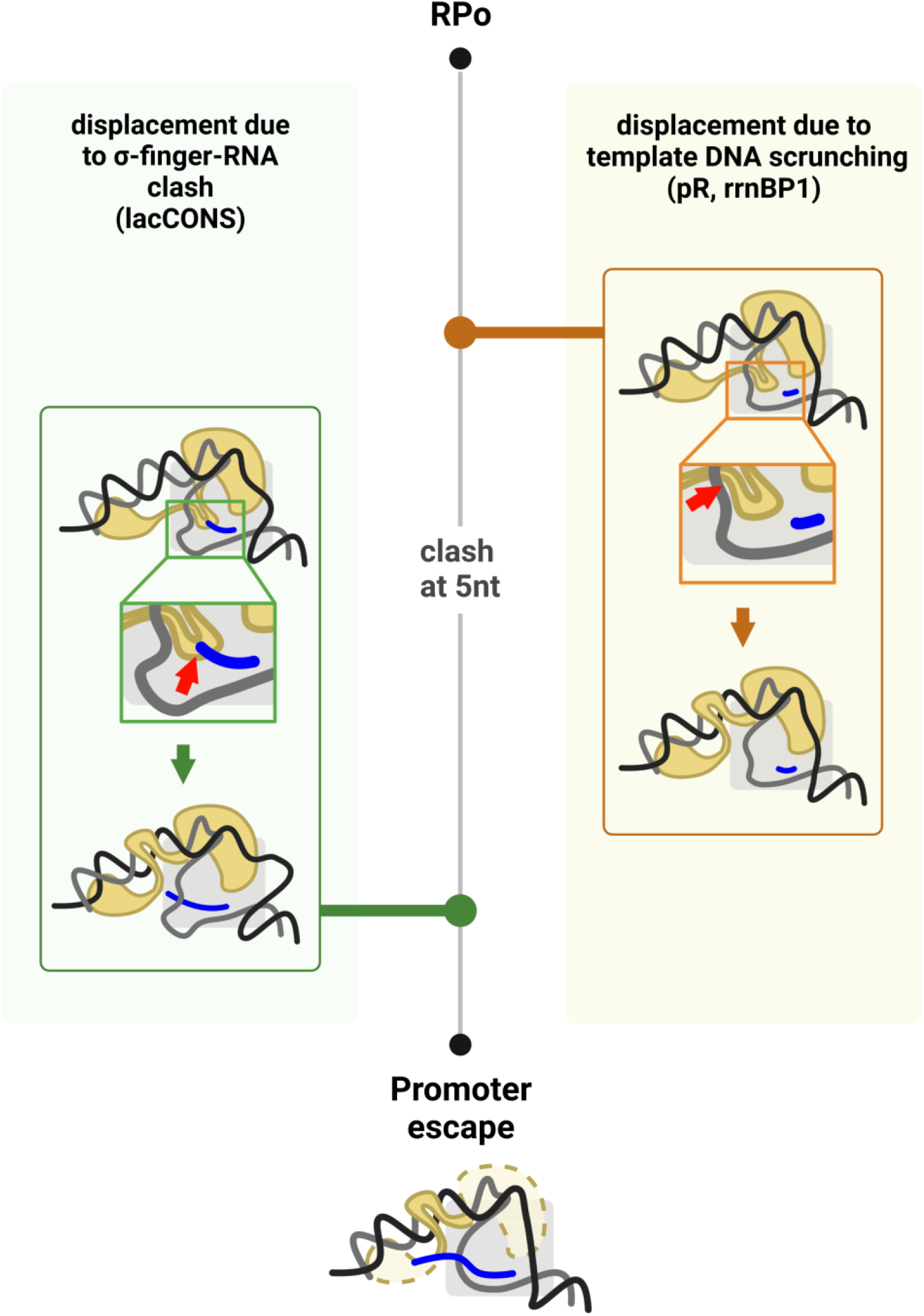
Two distinct, promoter-dependent mechanisms for the displacement of the σ-finger during initial transcription. ***Left***, displacement occurs due to the clash of the 5’-end of the RNA with the σ-finger. ***Right***, displacement occurs due to altered interactions of the finger with template DNA driven (directly or indirectly) by DNA scrunching.

### σ-finger displacement during initial transcription: a ribosomal RNA promoter (rrnBP1)

We then analysed σ-finger displacement on a ribosomal RNA promoter (rrnBP1); such ribosomal RNA promoters are responsible for a large fraction of transcription in bacteria, especially during fast growth; as such, these promoters are characterized by very high turnover rates. In contrast to pR, the rrnBP1 promoter is characterised by a very unstable RP_o_ (lifetime ∼ 1 s; Ref. 36) containing a 15-bp unwound bubble and a transcription start-site located nine bases downstream of the −10 element; this unusually long spacing (which is 2-nt longer than average) is associated with accumulation of some DNA scrunching in the open complex prior to RNA synthesis initiation^36^. Further, the rrnBP1 promoter is not associated with the production of significant amounts of abortive RNAs^36^, and it has been postulated that promoter escape on rrnBP1 may occur as early as RNA lengths of 3 nt^34^, although there is no direct physical evidence supporting this hypothesis.

Using the same approaches as for lacCONS and pR promoters, we observed that an RP_o_ formed on a rrnBP1 promoter fragment adopts almost exclusively (94% of molecules) the in-cleft σ-finger conformation, with the rest of the molecules adopting a displaced σ-finger conformation; these results are essentially identical to the profile observed for the lacCONS promoter. In sharp contrast, measurements performed in complexes able to synthesize increasing lengths of RNA showed a large increase in the fraction of displaced conformation as early as RP_itc≤2_ (29%; Fig. 4D). Strikingly, when dinucleotides used to form RP_itc≤2_ complexes were washed out from solution-- which should lead to dissociation of the RNA dinucleotide and re-formation of the initial RP_o_ -- the E* histogram still showed a significant fraction of complexes with a displaced σ-finger (24%; Fig. 4E). These results strongly suggest that, on rrnBP1, the formation of the first phosphodiester bond during the synthesis of the dinucleotide provided sufficient interactions to reposition the σ finger in a stably displaced conformation.

To define the point of escape for rrnBP1 and compare it to the point of σ-finger displacement, we designed a construct with donor and acceptor probes placed at an upstream (−15 of non-template strand) and a downstream (+20 of template strand) position of the rrnBP1 promoter (−15/+20 rrnBP1; Table S1). smFRET measurements on transcription complexes formed with the −15/+20 rrnBP1 (RP_o_, RP_itc≤2,_ RP_itc≤3,_ RP_itc≤4,_ and RP_itc≤5_) showed subpopulations with low FRET (E*∼0.15-0.20), which we assigned to conformations with no unwound promoter bubble, and subpopulations with intermediate FRET (E*∼0.35-0.40), which we assigned to conformations with unwound promoter bubbles in initial transcription (Fig. S13). Our results showed large subpopulations (88-95%) with intermediate FRET for RP_itc≤2,_ RP_itc≤3,_ and RP_itc≤4,_ indicating presence of unwound promoter bubbles characteristic of scrunched bubbles in initial transcription (Fig. S13A). Further extension of RNA to 5-mer (RP_itc≤5_) resulted in large decrease in the intermediate-FRET subpopulation, strongly suggesting that this length associates with loss of the scrunched unwound bubble, an event characteristic of promoter escape (Fig. S13A). We thus conclude that promoter escape in rrnBP1 occurs when a 5-nt RNA is synthesized. Taken together, our results for the rrnBP1 promoter indicate that the σ-finger is displaced well before promoter escape (at 5-nt RNA) and as early as the synthesis of a 2-nt RNA.

Real-time experiments with rrnBP1 promoter revealed two subpopulations with displacement events of ∼3.8 and ∼36 s, and E* efficiencies of ∼0.49 before NTP addition (Fig. 5D-E). The subpopulation exhibiting very fast displacement events was similar in abundance to subpopulations exhibiting fast displacement kinetics for the other two promoters (84% for rrnBP1 vs 88% for lacCONS and pR) but exhibited much faster displacement times (∼3.8 s for rrnBP1 vs. ∼14 s for lacCONS and ∼17 s for pR). This striking difference is a likely consequence of the distinctly different nature of RPo formed at the rrnBP1 promoter and its very early point of displacement (after synthesis of 2-nt RNA). The slow-displacing subpopulation on rrnBP1 is also 5-fold faster compared to the slow-displacing population in lacCONS (∼188 s) and pR (∼185 s) (Figs. 3C and 5B).

Overall, the results of the real time experiments reveal subpopulations of molecules showing very slow displacement events of ∼180-200 s for lacCONS and pR. Given the distinctly different point of displacement of the σ-finger for lacCONS (very late displacement) and for pR (early displacement) we suggest that the majority of the displacement time is spent before initial transcription begins. A likely explanation is that the slow displacers represent RPo molecules residing in a transcriptionally inactive conformation for a significant amount of time. Similar functional heterogeneity among RPo molecules has been recently reported in studies employing the pR promoter^37^ and the lacCONS promoter^38^. A subpopulation with such markedly slow displacement (∼180 s) is absent on rrnBP1. However, since we observe a very early displacement (after 2 nt RNA synthesis) for rrnBP1, it is unlikely the subpopulation with ∼36 s displacement time spent most of this time in initial transcription. Therefore, we suggest that, similar to the other two promoters, a long-lived transcriptionally inactive RP_o_ state also occurs on rrnBP1.

## DISCUSSION

### The σ-finger is displaced from an in-cleft conformational state during initial transcription

In this study, we developed a new FRET-based approach for detecting σ-finger conformations in single transcription complexes and captured the σ-finger conformational profile throughout initial transcription (from RP_o_ complexes to early elongation complexes) for three promoters with different transcription initiation profiles.

Notably, our FRET construct reports on the position of the base of the σ-finger (since we label residue 511) and cannot observe any motions that involve only the tip of the finger (residues 513-519), such as the motions observed in a structural study examining analogues of initial transcribing complexes with increasing RNA lengths^18^ (Fig. S1B). Therefore, the large FRET changes observed in our assays with different promoter fragments report on full displacement of the σ-finger from “in-cleft” conformations in early stages of transcription initiation. The observed FRET decrease upon RNA synthesis suggests that the σ-finger moves out of the RNAP main cleft and away from the leading edge of RNAP (where the other FRET probe is attached), likely moving towards the dsDNA upstream of the transcription bubble. Such displaced conformations have been proposed but were never directly observed in structural studies, likely due to high mobility of this displaced structural module.

Importantly, for all promoters studied, σ-finger displacement occurred at RNA lengths shorter than those required for promoter escape. Specifically, σ-finger displacements were observed after synthesis of 10-nt, 3-nt, and 2-nt RNA for the lacCONS, pR and rrnBP1 promoters, respectively, whereas promoter escape occurred after synthesis of 12-nt, 11-nt and 5-nt RNA, respectively (based on results in Fig. S7, Henderson et al.^34^ and Fig. S13).

### σ-finger displacement can occur via different mechanisms depending on the promoter

Structural work on the σ^70^ finger using synthetic transcription initiation scaffolds containing different lengths of RNA revealed that, as the RNA length increases, the 5’-end of the RNA clashes with the σ-finger; and for sufficiently long RNAs, the tip of the σ-finger gets displaced, folding back on itself in a manner analogous to compression of a spring, with each stepwise increase in RNA-length leading to greater displacement of the tip of the finger^18^. Very recent structural work on initial transcribing complexes formed by σ^54^ RNAP and a synthetic promoter scaffold containing increasing lengths of pre-synthesised RNA showed that the “RII-finger”, a σ^54^ element functionally equivalent to the σ^70^ finger, also folds back on itself when the RNA reaches ∼7 nt, at which point a steric clash between the RNA and the RII-finger occurs in a manner analogous to the σ^70^-finger ^39^. These studies support the hypothesis that full displacement of the σ-finger away from the path of the extending RNA follows an obligatory steric clash with the extending RNA. As a corollary of this “obligatory steric clash” hypothesis, the exact point of displacement for different promoters should be dictated by the extent to which the σ-finger protrudes inside the unwound transcription bubble and the strength of the contacts it makes with the template ssDNA and the RNA.

Consistent with the obligatory steric clash hypothesis, σ-finger displacement from the cleft occurred late on the lacCONS promoter -- after synthesis of 10-nt RNA. Intriguingly, however, and contrary to the obligatory steric clash hypothesis, our measurements with the pR and rrnBP1 promoters showed that displacement occurred at RNA lengths of only 3-nt and 2-nt, respectively, lengths much shorter than the 5-nt RNA length expected for a steric clash of the 5’-end of the RNA with the σ-finger for these promoters (Figs. 4C-D). We thus suggest that, even before the ensuing steric clash between the σ-finger and the nascent RNA in the active-site cleft of RNAP, a steric clash between the σ-finger and the scrunched template DNA, and/or a weakening of interactions between the σ-finger and the template ssDNA due to scrunching, triggers disruption of the σ-finger contacts, enabling full displacement of the σ-finger. Notably, the σ-finger interacts with template bases −3/-4 with promoters of a canonical discriminator of 6-nt (such as lacCONS and λPr [Ref. 7]), and template bases −4/-5 with promoters with a longer discriminator (such as *rrn*BP1 [Ref. 40]); the template bases involved in these contacts are different in all 3 promoters, raising the possibility of a DNA-sequence determinant for finger displacement. Further, the recent work on the σ^54^ RII-finger displacement also showed altered interactions between the RII-finger and the template DNA due to DNA scrunching^39^.

### Influence of the different mechanisms of σ-finger displacement on the “initiation pause”

Previous single-molecule and crosslinking studies of initial transcription with the lacCONS promoter identified an “initiation pause” during extension of a 6-nt to a 7-nt RNA; the same study showed that deletion of the σ-finger resulted in significantly reduced pause lifetimes as RNA extends from 6-nt to 7-nt, establishing the σ-finger as a crucial determinant of pausing during initial transcription, in addition to the initial transcribed sequence^23, 31, 33^. We suggest that, for lacCONS, the σ-finger clashes with the RNA after synthesis of a 5-nt RNA, while making additional stabilising contacts with the 5’-end of the RNA, as observed in Li et al.^18^, likely raising the energy barrier of translocation during initial transcription and enhancing pausing, as observed^23, 41^. Further extension of RNA beyond the 7-mer results in additional scrunched DNA and additional folding of the tip of the σ-finger, as observed in Li et al.^18^ – both structural elements capable of acting like a compressed spring and storing energy which can be released to disrupt RNAP promoter contacts during late stages of initial transcription, eventually enabling escape from the promoter. Our observation of the σ-finger getting displaced at 10-nt RNA length – well before the point of promoter escape for lacCONS (Fig. S7) suggests the stored energy in the folded σ-finger may indeed be used up in this step, disrupting the contacts formed between the σ-finger and the RNA, facilitating its displacement from the active centre cleft, and clearing the path for promoter escape.

However, for the pR and rrnBP1 promoters, an early displacement of the σ-finger well before its expected clash with the nascent RNA chain indicates that the additional translocation energy barrier posed by the σ-finger (as e.g., in the case of lacCONS) would be absent. Hence, one expects significantly reduced lifetimes of the “initiation pause” for these promoters, leading to much shorter times spent in initial transcription. We thus suggest that the σ-finger and its displacement can act as a major determinant of rates of transcription initiation and promoter escape.

### Interactions with the 5’-end of nascent RNA modulate σ-finger displacement

In contrast to our results obtained for primer-dependent transcription initiation, primer-independent transcription initiation yielded strikingly different results for the lacCONS promoter, with σ-finger displacements observed at a much earlier stage (after synthesis of 5-nt RNA instead of 10-nt RNA). Primer-dependent transcription involves synthesis of an RNA with a hydroxyl group at the 5’-end (5’-OH), while primer-independent transcription involves an RNA with a triphosphate at the 5’-end (5’-ppp). Since the bacterial σ-finger is negatively charged under physiological conditions (due to the presence of a stretch of four acidic residues in the σ-finger: D513, D514, E515, and D516), our results indicates that the bulky, negatively-charged triphosphate group at the 5’-end of the growing RNA repels the negatively charged σ-finger, triggering σ-finger displacement at RNA lengths significantly shorter than that observed for reactions involving 5’-OH ends of the RNA. As argued in the previous section, a displacement after 5-nt RNA synthesis would result in a lower energy barrier of translocation for the initiation pause (observed at +6) for lacCONS and consequently one expects a much shorter pause lifetime in case of primer-independent initial transcription for this promoter. In agreement with this result, Dulin et al.^31^, previously reported a ∼2.5-fold decrease in pause duration for primer-independent initiation as compared to primer-dependent initiation at the same promoter^30^.

Notably, most transcription under exponential growth involve primer-independent initiation, while primer-dependent initiation becomes prevalent during stationary phase^42, 43^. Our results indicate that σ-finger interactions with the 5’-end of the RNA may be used by bacteria to tune gene expression in a growth-dependent fashion, since an early σ-finger displacement in primer-independent initiation could facilitate promoter escape and result in higher transcription output during exponential growth, while larger timescales for σ-finger displacement in primer-dependent initiation, as shown by real-time measurements, could lead to reduced transcription during stationary phase.

### σ-finger displacement on a single promoter occurs at substantially different timescales

Our real-time measurements of displacement time for primer-dependent initiation on lacCONS revealed that only ∼25% of the molecules show displacement. Since we miss the displacement event in ∼20% of the molecules (due to probe photobleaching prematurely terminating our FRET time-trajectories), our results imply that ∼55% of the molecules are either transcriptionally inactive or remain in early stages of initial transcription where the σ-finger is still not displaced. These results are consistent with previous reports from biochemical and single-molecule experiments showing the presence of a class of transcription complexes that remain in abortive initiation and do not proceed to elongation^31, 34^.

The molecules showing displacement belonged to two sub-classes, exhibiting either a short (5 – 20 s) or much longer (>150 s) displacement time for the lacCONS promoter. These times are strikingly similar to the promoter-escape times [∼13 s (96%) and ∼200 s (4%)] observed in a single-molecule study on the same promoter^38^. Further analysis revealed σ-finger conformations with distinct mean E* values in the initial state (i.e., in the RP_o_ state) for the kinetically different molecules, indicating that the heterogeneity in displacement times is correlated with conformationally different RP_o_ molecules. How do molecules with different σ-finger positions result in different displacement kinetics? We envision that the basis for this resides on altered pre-organisation of the template strand by the differently positioned σ-fingers, and/or on differences in the ensuing steric clash between the nascent RNA and the differently positioned σ-finger modules.

In contrast, our analysis on the pR promoter (which showed displacement early in initial transcription, after synthesis of a 3-mer RNA) showed both short and long displacement times, as in the lacCONS case – however, both short and long displacement times show σ-finger conformations with similar E* values before NTP addition. What gives rises to this kinetic heterogeneity? Given the short RNA length needed for σ-finger displacement on the pR promoter, a steric clash with the RNA during initial transcription can be ruled out; further, our assay could not pick up significantly different σ-finger conformations in the kinetically distinct subpopulations and, as such, our results on λP_R_ do not support the presence of an altered pre-organisation of the template strand driven by different σ-finger conformations.

Instead, we speculate that the kinetic heterogeneity in this case arises from conformational heterogeneity in RP_o_ molecules which involve different template strand organizations as observed in a recent study on RP_o_ formed on a pR promoter^19^; in that study, the authors report one RP_o_ conformation with template strand conformations incompatible with binding initiating nucleotides, and hence incompatible with transcription initiation^19^. In this scenario, the long σ-finger displacement times for a subpopulation of transcription complexes on the pR promoter could originate from the rate-limiting step of the conversion of the inactive conformation of RP_o_ to the active conformation.

Our analysis on the rrnBP1 promoter also showed kinetically heterogeneous σ-finger displacement, with both fast-displacers and slow-displacers showing displacement much faster than the other two promoters. This heterogeneity is intriguing, since the displacement times for rrnBP1 under our conditions essentially report on the binding of the dinucleotide primer (and not on rate of RNA synthesis); in other words, primer binding alone can trigger σ-finger displacement. The observed heterogeneity again indicates the presence of distinct RP_o_ conformational states (as for the pR promoter), with different template strand organisation, which in turn result in distinct binding efficiency of the dinucleotide primer to the RNAP active site. Similar to the pR promoter, we do not observe significant differences in E* values for the σ-finger conformation for the kinetically different molecules.

Taken together, our study identifies displacement of the σ-finger as a key step that influence the kinetics of initial transcription and reveals different mechanisms which influence this process; these mechanisms can be leveraged by the bacteria to regulate transcription at many levels (different genes, different sets of genes, or different physiological states). Such conformational control may also offer a new target for generating antibiotics, e.g., by identifying small molecules that severely delay or block promoter escape on pivotal genes, such as the rRNA genes.

We note that structural modules similar the bacterial σ-finger (in that they protrude into the cleft of transcription initiation complexes, interact with template ssDNA and occupy the path of the nascent RNA chain) are present in other kingdoms of life [e.g., TFB zinc ribbon and CSB of the archaeal RNAP^10^; Rrn7 zinc ribbon and B reader R of RNAP-I^14, 15^; TFIIB zinc ribbon and B reader of RNAP-II^11, 13, 44, 45^; and Brf1 zinc ribbon of RNAP-III^16, 17^. We suggest these modules likely behave in a manner analogous to the bacterial σ-finger and influence the kinetics of transcription and gene regulation in both archaea and higher eukaryotes in an analogous manner.

## Materials and Methods

### Accessible volume calculations

Accessible volume calculations for DL RNAP-σ^70^ (labelled at σ^70^ residue 366 Ser and residue 511 Ile) were carried on crystal structures PDB 4G7H, 6KQD, 6KQE, 6KQF, 6KQG, 6KQH, 4YLN, 7KHB, 7MKD and 7MKE using FPS software^26^. Amino acid side chains on σ^70^ residues 511 and 366 (4YLN, 7KHB, 7MKD and 7MKE), on σ^A^ residues 321 and 174 (6KQD, 6KQE and 6KQF) and on σ^A^ residues 319 and 174 (4G7H, 6KQD, 6KQE, 6KQF, 6KQG and 6KQH) were deleted in all structures. The dye attachment point used was Cα. Dye parameters for accessible volume measurements were as follows: Alexa647 maleimide (linker length = 21Å, linker width = 4.5Å and dye radii = 11.0Å, 4.7Å, 1.5Å) and Cy3B maleimide (linker length = 9.1Å, linker width = 4.5Å and dye radii = 7.7Å, 2.5Å, 1.3Å). Average distances and corresponding FRET values were then calculated using a Förster radius of 60Å.

### Preparation of reagents

#### σ^70^ derivatives

Double cysteine modified σ^70^ derivatives were prepared as follows: single colonies of *E. coli* strain BL21(DE3) (Millipore) were co-transformed with a pGEMD (-Cys) derivative encoding two cysteine residues at positions 366 and 511 [constructed from plasmid pGEMD (-Cys)^46^ by use of site-directed mutagenesis (QuikChange Site-Directed Mutagenesis Kit; Agilent) to replace codons 366 and 511 by a codon encoding cysteine residue], were used to inoculate 20 ml LB broth containing 100 μg/ml ampicillin, and cultures were incubated 16 h at 37°C with shaking. Culture aliquots (2x10 ml) were used to inoculate LB broth (2x1 L) containing 100 μg/ml ampicillin; cultures were incubated at 37°C with shaking until OD_600_ = 0.7; IPTG was added to 1 mM; and cultures were further incubated for 4 h at 37°C with shaking. Cells were harvested by centrifugation (5,000 x g; 20 min at 4°C), re-suspended in 50 ml lysis buffer [40 m*M* Tris–HCl (pH 7.9), 300 m*M* NaCl, 1 m*M* EDTA, one protease inhibitor cocktail tablet, and 0.2% deoxycholate], and lysed by emulsification (Emulsiflex-C5; Avestin, Inc., Ottawa, Canada). Inclusion bodies containing σ^70^ derivatives were isolated by centrifugation (10,000 *g*; 20 min at 4°C), washed with 20 ml lysis buffer containing 0.2 mg/ml lysozyme and 0.5% Triton X-100, and washed with 20 ml lysis buffer containing 0.5% Triton X-100 and 1 m*M* DTT [with each wash step involving sonication 2 × 1 min at 4 ° in wash buffer, incubation 10 min at 4°C in wash buffer, and centrifugation (10,000 *g*; 20 min at 4 °)]. Washed inclusion bodies containing σ^70^ derivatives were solubilized in 40 ml 6 *M* guanidine–HCl, 50 m*M* Tris–HCl (pH 7.9), 10 m*M* MgCl_2_, 10 μ*M* ZnCl_2_, 1 m*M* EDTA, 10 m*M* DTT, and 10% glycerol, and dialyzed against 2 L TGED [20 m*M* Tris–HCl (pH 7.9), 0.1 m*M* EDTA, 0.1 m*M* DTT, and 5% glycerol] containing 0.2 *M* NaCl (20 h at 4 °; two changes of buffer). The sample was centrifuged (10,000 *g*; 20 min at 4 °) to remove particulates and applied to a Mono-Q HR 10/10 column (Amersham-Pharmacia Biotech, Piscataway, NJ) pre-equilibrated in the same buffer. The column was washed with 16 ml of the pre-equilibration buffer and eluted in 2-ml fractions of a 160-ml linear gradient of 200–600 m*M* NaCl in TGED (with σ^70^ derivatives typically eluting at ∼360 m*M* NaCl in TGED). Fractions containing the σ^70^ derivative were identified by SDS–PAGE and Coomassie staining and are pooled. Pooled fractions were concentrated in storage buffer [20 mM Tris-HCl, pH 7.9, 200 mM NaCl, 0.1 mM EDTA, 1 mM TCEP, and 5% glycerol] using 10 kDa MWCO Amicon Ultra-15 centrifugal ultrafilters, and stored in aliquots at −80°C.

Fluorescent-probe-labelled, σ^70^ derivatives were prepared as follows: A reaction mixture containing 10 μM σ70 derivative [with cysteines at position 366 and 511], 500 μM Cy3B maleimide and 500 μM Alexa647 maleimide in 0.5 ml buffer B (100 mM potassium phosphate buffer pH 8.0, 50 mM NaCl, 1 mM EDTA, 5 mM TCEP, and 2% dimethylformamide) was incubated 4 h on ice, subjected to 5 cycles of buffer exchange (dilution with 5 ml buffer B (without dimethyl formamide), followed by concentration to 0.5 ml using 10 kDa MWCO Amicon Ultra-15 centrifugal ultrafilters (EMD Millipore), and stored in aliquots at −80°C.

Efficiencies of incorporation of fluorescent probes were determined from UV/Vis-absorbance measurements and were calculated as:

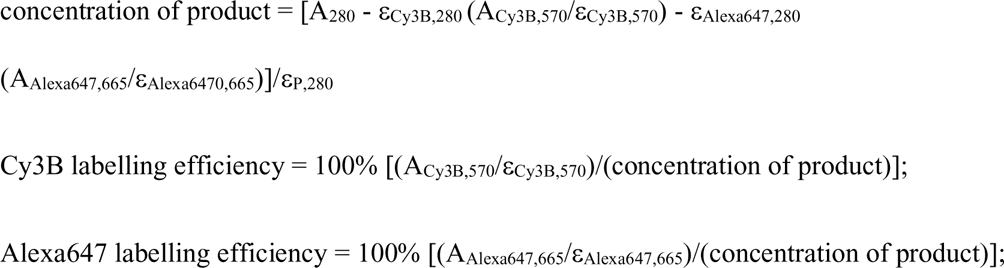

where A_280_ is the measured absorbance at 280 nm, A_Cy3B,570_ is the measured absorbance of Cy3B (570 nm), A_Alexa647,665_ is the measured absorbance of Alexa647 (665 nm), ε_P,280_ is the molar extinction coefficient of σ70 at 280 nm (39,760 M^−1^ cm^−1^), ε_Cy3B,280_ is the molar extinction coefficient of Cy3B (10,400 M^−1^ cm^−1^) at 280 nm, ε_Alexa647,280_ is the molar extinction coefficient of Alexa647 at 280 nm (7,350 M^−1^ cm^−1^), ε_Cy3B,570_ is the extinction coefficient of Cy3B at 570 nm (130,000 M^−1^ cm^−1^), and ε_Alexa647,665_ is the extinction coefficient of Alexa647 at 665 nm (245,000 M^−1^ cm^−1^).

#### RNAP core enzyme

Hexahistidine-tagged *Escherichia coli* RNAP core enzyme was prepared using co-expressed genes encoding RNAP β’, β, α, and ω subunits to afford an RNAP core enzyme as follows: single colonies of *E. coli* strain BL21(DE3) (Millipore) co-transformed with plasmid pEcABC-His_6_^47^ and plasmid pCDFω^48^, were used to inoculate 20 ml LB broth containing 100 μg/ml ampicillin, and 50 μg/ml kanamycin. Cultures were incubated for 16 h at 37°C with shaking. Culture aliquots (2x10 ml) were used to inoculate LB broth (2x1 L) containing 100 μg/ml ampicillin, and 50 μg/ml kanamycin; cultures were incubated at 37°C with shaking until OD_600_ = 0.6; and IPTG was added to 1 mM; and cultures were further incubated 16 h at 16°C with shaking. Cells were harvested by centrifugation (4,000 x g; 20 min at 4°C), re-suspended in 20 ml buffer C (10 mM Tris-HCl, pH 7.9, 200 mM NaCl, and 5% glycerol), and lysed using an EmulsiFlex-C5 cell disrupter (Avestin). The lysate was cleared by centrifugation (20,000 x g; 30 min at 4°C), precipitated with polyethyleneimine (Sigma-Aldrich) as in Niu et al.^49^, and precipitated with ammonium sulfate as in Niu et al.^49^.The precipitate was dissolved in 30 ml buffer C and loaded onto a 5 ml column of Ni-NTA-agarose (Qiagen) pre-equilibrated in buffer C, and the column was washed with 50 ml buffer C containing 10 mM imidazole and eluted with 25 ml buffer C containing 200 mM imidazole. The sample was further purified by anion-exchange chromatography on Mono Q 10/100 GL (GE Healthcare; 160 ml linear gradient of 300-500 mM NaCl in 10 mM Tris-HCl, pH 7.9, 0.1 mM EDTA, and 5% glycerol; flow rate = 2 ml/min). Fractions containing hexahistidine-tagged *E. coli* RNAP core enzyme were pooled, concentrated to ∼1 mg/ml using 30 kDa MWCO Amicon Ultra-15 centrifugal ultrafilters (EMD Millipore), and stored in aliquots at −80°C.

#### DL RNAP-σ^70^

DL RNAP-σ^70^ was formed by incubating hexahistidine tagged RNAP core enzyme with doubly labelled σ^70^ derivative in the ratio 1:2 in KG7 buffer (40 mM HEPES-NaOH, pH 7.0, 100 mM potassium glutamate, 10 mM MgCl_2_, 1 mM dithiothreitol and 5% glycerol) at 30°C for 30 min, and then on ice for 12 hrs.

#### Nucleic acids

Lyophilized oligodeoxyribonucleotides (BiomersGmBH and IDT) were dissolved in nuclease-free water (Ambion) to a final concentration of 100mM and stored at −20°C. For experiments involving DL RNAP-σ^70^, non-template oligodeoxyribonucleotides contained a 5’-biotinylated end. For experiments involving fluorescent-labelled oligodeoxyribonucleotides, non-template oligodeoxyribonucleotides were labelled with Cy3B and template oligodeoxyribonucleotides with Alexa647 in the positions stated (Table S1).

Double-stranded DNA was prepared as follows: non-template oligodeoxyribonucleotides (1 μM) and template oligodeoxyribonucleotides (1.1 μM for annealing with biotinylated oligodeoxyribonucleotides and 1 μM for annealing with nonbiotinylated oligodeoxyribonucleotides) in 50 μl 10mM Tris-HCl, pH 7.9 and 0.2M NaCl were heated for 5 min at 95°C and then cooled to 25°C in 2°C steps with 1 min per step in a thermal cycler (Applied Biosystems). The resulting double-stranded DNA (∼1 μM) was stored at −20°C.

#### Small molecules

NTPs (New England Biolabs) and dinucleotides (Trilink, Biolog LSI GmbH) were diluted in nuclease-free water (Ambion) to a final concentration of 100 mM or 10 mM and stored at −20°C.

#### RP_o_

For experiments involving DL RNAP-σ^70^, RP_o_ was formed as follows: 15 nM DL RNAP-σ^70^ was incubated with 10 nM biotinylated double-stranded promoter DNA in 10 μl KG7 (40 mM HEPES-NaOH, pH 7.0, 100 mM potassium glutamate, 10 mM MgCl_2_, 1 mM dithiothreitol and 5% glycerol) at 37°C for 20 min. For experiments involving double fluorescent-labelled dsDNA, RP_o_ concentrations of 10 nM hexahistidine-tagged RNAP holoenzyme and 30 nM double-labelled dsDNA were used.

#### smFRET using TIRF-ALEX: sample preparation

Observation chambers with biotin-PEG-passivated glass floors were functionalized with neutravidin (Sigma Aldrich). For experiments involving double-labelled dsDNA, the glass was additionally treated with biotinylated anti-hexahistidine monoclonal antibody (Penta-His Biotin Conjugate; Qiagen) as in Duchi et al.^23^ and Dulin et al.^31^.

For experiments involving DL RNAP-σ^70^ in Figs. 1-5, S5, S6, S8, S9, S10 and S12, RP_o_ complexes containing biotinylated non-template strands were immobilized as follows: 50 μl of 0.1 nM double fluorescent-labelled RP_o_ in KG7 were added to the observation chamber and incubated for 10-30 s at 22°C. Wells were then washed with 3x50 μl KG7. For experiments involving double labelled dsDNA and the DL RNAP-σ^70^ holoenzyme (Figs. S4, S7 and S13), complexes containing hexahistidine tags were immobilised in a similar fashion but incubated for 2-4 min in observation chambers which had been additionally treated with biotinylated anti-hexahistidine monoclonal antibody.

For experiments in Figs. 2 (except RP_itc2_), 4 (except RP_o_ and RP_itc2_), S5, S7 and S13, preformed RP_o_ were immobilised as described previously, and a 50 μl solution of the specified combination of NTPs (100 μM) and dinucleotides (500 μM) in KG7 were added and incubated in the observation well for 5 min at 22°C. Observation wells were then washed with 5x50 μl KG7 and 50 μl imaging buffer (KG7 containing 2 mM Trolox (Sigma-Aldrich), 1 mg/ml glucose oxidase (Sigma-Aldrich), 40 μg/ml catalase (from bovine liver C30; Sigma-Aldrich), and 8 mM D-glucose) added. Data were then acquired. For RP_o_ experiments in Figs. 4 and S12, preformed RP_o_ were immobilised; subsequently, 50 μl of imaging buffer was added to the well, and the well was imaged. For RP_itc2_ experiments in Figs. 1D, 2 (RP_itc2_), 4(RP_itc2_), S6 and S13, preformed RP_o_ were immobilised and 50 μl imaging buffer supplemented with the 500 μM of the relevant dinucleotide and imaged. For the experiment in Fig. 4E, preformed RP_o_ was immobilised and 500 μM ApC added to the well and incubated for 5 min. The well was then washed with 5x50 μl KG7, 50 μl imaging buffer added and the well was imaged.

For real-time experiments with the lacCONS promoter DNA in Figs. 3, S8, S9 and S10, RP_o_ were prepared as before and 40 μl of imaging buffer supplemented with 625 μM dinucleotide (ApA or pppApA) was added to the well and incubated for 2 min at 22°C. Data acquisition was started and a 10 μl aliquot of 5 mM GTP/UTP/ATP in imaging buffer was added into the well and mixed. Final concentrations were 500 μM for the dinucleotide and 1 mM for the NTPs.

For real-time experiments involving the pR promoter in Fig. 5A-C, experiments were performed in a similar fashion with RP_o_ complexes prepared as above. An aliquot of 37.5 μl imaging buffer was added to the well. Data acquisition was then started and a 12.5 μl aliquot of 2 mM ApU, 4 mM GTP/UTP/ATP added into the well and mixed. Final concentrations were 500 μM of ApU and 1 mM NTPs. The same method was used for real-time experiments involving the rrnBP1 promoter in Fig. 5D-F but with final concentration of 500 μM ApC and 1 mM GTP/UTP/ATP/CTP instead.

For experiments in Fig. S7, RP_o_ complexes were prepared as above, and a 50 μl solution of 500 μM ApA and 100 μM UTP/GTP/ATP in KG7 was incubated in the well for 5 min at 22°C to form RP_itc11_. Wells were washed with 5x50 μl KG7 and 50 μl imaging buffer added. Data were then acquired for RP_itc11_. Immediately after the imaging, wells were washed with 5x50 μl KG7, and a 50 μl aliquot of 1 mM GTP/CTP was incubated into the well for 5 min at 22°C to form RD_e14_. Data was then acquired for RD_e14_.

#### smFRET using TIRF-ALEX: data collection

smFRET experimental data was collected using a custom-built objective-type TIRF microscope. Light from green (532 nm, Cobolt) and red (635 nm, Coherent) lasers were combined using a dichroic mirror and coupled into single-mode fiber. The output was focused onto the back focal plane of a 100x oil-immersion objective (NA 1.4, Olympus) and laterally displaced of the optical axis such that the incident angle at the oil-glass interface of the observation chamber exceeded the critical angle and created an evanescent illumination wave. Alternating-laser excitation (ALEX) was achieved through an acousto-optical modulator. A dichroic mirror (545nm/650nm, Semrock) and emission filters (545nm LP Chroma and 633/25nm notch, Semrock) separated the fluorescence emission from the excitation. The fluorescence emission was then focussed on a slit and spectrally separated using a dichroic mirror (630nm DLRP, Omega) into two channels focussed onto an EMCCD (iXon 897, Andor). A motorised x/y-scanning stage (MS-2000, ASI) was used to mount the observation chambers.

All data acquisition was carried out at 22°C using ALEX-TIRF. For experiments in Figs. 1C, 2A-B and 4 was collected for 50 s at a frame rate of 100 ms/frame (50 ms ALEX) using laser powers of 0.7 mW (532 nm laser) and 0.3 mW (635 nm laser). For experiments in Fig. S7 and S13, data were collected for 50 s at a frame rate of 100 ms/frame (50 ms ALEX) using laser power of 0.6 mW (532 nm laser) and 0.2 mW (635 nm laser). For experiments in Fig. 3 and 5, data was collected for 500 s at a frame rate of 400 ms/frame.

#### smFRET using TIRF-ALEX: data collection

Background corrected fluorescence-emission intensity vs time trajectories for donor emission upon donor excitation (I_DD_), acceptor emission upon donor excitation (I_DA_) and acceptor emission upon acceptor excitation (I_AA_) were extracted using software package Twotone-ALEX^50^. Intensity vs time trajectories were manually inspected to exclude trajectories exhibiting I_DD_<100 or >1500 counts, I_AA_<100 or >1000 counts, multi-step donor or acceptor photobleaching and donor or acceptor photoblinking. Selected trajectories were then used to calculate trajectories of apparent FRET efficiencies (E*) and donor-acceptor stoichiometry (S) using equations:

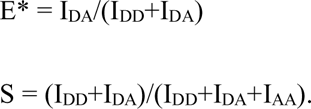

S values were used to distinguish between species containing both donor and acceptor fluorophores and species containing only one of either donor or acceptor fluorophore. E* histograms for species containing both donor and acceptor fluorophores were built and fitted to Gaussian distributions using Origin (Origin Lab) to provide subpopulation percentages and mean E* values as in Figs. 1D, 2, 3E, 4C-E, 5C, 5F, S4 and S12. In Fig. 2C, the error in the low E* “displaced” subpopulation percentage was calculated as follows; datasets from experiments in Figs. 2A and 2B were split into different their individual experimental datasets, where it had been performed more than once (or into three subsets, where the experiment was only performed once). These data subsets were then individually plotted and fitted to Gaussian distributions to find a mean and standard deviation in the mean of the low-E*“displaced” subpopulation percentage as plotted in Fig. 2C.

To identify any dynamics and their time scales, E*-time trajectories were analysed using Hidden Markov Modelling (HMM) as implemented in the software package ebFRET (Ref. 30). By maximizing the lower bound, a three-state model was chosen to fit any dynamic E*-time trajectories in experiments involving DL RNAP-σ^70^. For experiments in Figs. 1E, 2, S4, S5 and S6, dynamic molecules were defined as those with three or more transitions between different states. Fitted E* values from the trajectories classified into the three-state model were then plotted and fitted to Gaussian distributions in Origin to extract population percentages and a mean E* value for each state. Dwell times were extracted and plotted as histograms in S4, S5 and S6, which were then fit to single exponential decays in all cases, apart from the histogram in S6 where a peaked exponential decay was used for the low-FRET state. To estimate the error in the percentage of dynamic molecules shown in Fig. 2D, datasets from experiments in Fig. 2A and 2B were split into different their individual experimental datasets, where it had been performed more than once (or into three subsets, where the experiment was only performed once). The percentage of dynamic molecules for each data subset was then individually calculated to find a mean and standard deviation in the mean of the % of dynamic molecules as plotted in Fig. 2D.

For real time experiments in Figs. 3, 5, S8, S9 and S10, all E*-time trajectories were analysed initially using HMM and then categorised into four groups as described in the main text. After categorization, HMM analysis was re-run on groups exhibiting displacement (4-state model to incorporate the displaced state) or dynamics (3-state model) separately. Displacement time histograms for σ-finger displacement (Figs. 3C, 5B, 5E, S10A) were calculated by subtracting the time of nucleotide addition from the time of σ-finger displacement, taken from HMM analysis. The time for σ-finger displacement were then plotted as histograms and fit to double exponential decays in Origin and the two time-constants extracted. For analysis in Figs. 3E, 5C, 5F and S10B, E* values before nucleotide addition were extracted from traces exhibiting σ-finger displacement (Class-I and Class-II). These traces were classified into either exhibiting ‘short’ or ‘long’ finger displacement using a boundary calculated from the time at which the two components of the double exponential decay intersect. The E*-values for these two groups were then extracted and a histogram produced from binned values. These were then fit to single gaussian distributions in Origin.

## ACKNOWLEDGEMENTS

We thank Konstantin Brodolin and Xiaodong Zhang for critical reading of the manuscript, and Zakia Morichaud for *in vitro* transcription assays. We also thank Xiaodong Zhang for sharing unpublished results from her group prior to publication. This work was supported by the Wellcome Trust (110164/Z/15/Z to A.N.K.), the UK BBSRC (BB/R008655/1 to A.N.K.), and a UK EPSRC studentship (project 2285264 to A.W.). Many elements in Figures were created using BioRender.

