## Supplemental table and figures for "The displacement of the σ^70^ finger in initial transcription is highly heterogeneous and promoter-dependent"

**SUPPLEMENTAL INFORMATION FOR WANG ET AL.**

**Table S1.** DNA sequences used.

| | |
|---|---|
| **lacCONS+12C** construct for measurements in Figs. 1D, 2A ($RP_{itc2}$, $RP_{itc4}$, $RP_{itc7}$, $RP_{itc11}$), 2B ($RP_{itc2}$, $RP_{itc4}$, $RP_{itc11}$), S5, S6, S12 | |
| Non-template strand | 5’-(biotin)AGGCTTGACACTTTATGCTTCGGCTCGTATA ATGTGTGGAATTGTGAGAGCGGATAACAATTTC-3’ |
| Template strand | 5- GAAATTGTTATCCGCTCTCACAATTCCACACATTAT ACGAGCCGAAGCATAAAGTGTCAAGCCT-3’ |
| **lacCONS+6A+10C** construct for measurements in Figs. 2A ($RP_{itc5}$, $RP_{itc9}$), 2B ($RP_{itc9}$) | |
| Non-template strand | 5’-(biotin)AGGCTTGACACTTTATGCTTCGGCTCGTATA ATGTGTGGAATTGAGAGCGCGGATAACAATTTC-3’ |
| Template strand | 5’- GAAATTGTTATCCGCGCTCTCAATTCCACACATTAT ACGAGCCGAAGCATAAAGTGTCAAGCCT-3’ |
| **lacCONS+6C** construct for measurements in Fig. 2B ($RP_{itc5}$) | |
| Non-template strand | 5’-(biotin)AGGCTTGACACTTTATGCTTCGGCTCGTATA ATGTGTGGAATTGCGAGAGCGGATAACAATTTC-3’ |
| Template strand | 5’- GAAATTGTTATCCGCTCTCGCAATTCCACACATTAT ACGAGCCGAAGCATAAAGTGTCAAGCCT -3’ |
| **lacCONS+7C** construct for measurements in Figs. 2A ($RP_{itc6}$), 2B ($RP_{itc6}$) | |
| Non-template strand | 5’-(biotin)AGGCTTGACACTTTATGCTTCGGCTCGTATA ATGTGTGGAATTGTCAGAGCGGATAACAATTTC-3’ |
| Template strand | 5’- GAAATTGTTATCCGCTCTGACAATTCCACACATTAT ACGAGCCGAAGCATAAAGTGTCAAGCCT-3’ |

| | |
|---|---|
| **lacCONS+8C** construct for measurements in Fig. 2B ($RP_{itc7}$) | |
| Non-template strand | 5'-(biotin)AGGCTTGACACTTTATGCTTCGGCTCGTATA ATGTGTGGAATTGTGCGAGCGGATAACAATTTC-3' |
| Template strand | 5'-GAAATTGTTATCCGCTCGCACAATTCCACACATTAT ACGAGCCGAAGCATAAAGTGTCAAGCCT-3' |
| **lacCONS+9C** construct for measurements in Figs. 2A ($RP_{itc8}$), 2B ($RP_{itc8}$) | |
| Non-template strand | 5'-(biotin)AGGCTTGACACTTTATGCTTCGGCTCGTATA ATGTGTGGAATTGTGACAGCGGATAACAATTTC-3' |
| Template strand | 5'- GAAATTGTTATCCGCTGTCACAATTCCACACATTAT ACGAGCCGAAGCATAAAGTGTCAAGCCT-3' |
| **lacCONS+11C** construct for measurements in Figs. 2A ($RP_{itc10}$), 2B ($RP_{itc10}$) | |
| Non-template strand | 5'-(biotin)AGGCTTGACACTTTATGCTTCGGCTCGTATA ATGTGTGGAATTGTGAGACCGGATAACAATTTC-3' |
| Template strand | 5'- GAAATTGTTATCCGGTCTCACAATTCCACACATTAT ACGAGCCGAAGCATAAAGTGTCAAGCCT-3' |
| **lacCONS+15C** construct for measurements in Figs. 2A ($RD_{e14}$), 2B ($RD_{e14}$), 3, S8, S9, S10 | |
| Non-template strand | 5'-(biotin)AGGCTTGACACTTTATGCTTCGGCTCGTATA ATGTGTGGAATTGTGAGGAGGACGGATAACAATTTC-3' |
| Template strand | 5'- GAAATTGTTATCCGTCCTCCTCACAATTCCACACAT TATACGAGCCGAAGCATAAAGTGTCAAGCCT-3' |
| **lacCONS -15/+20** construct for measurements in Fig. S7 | |

| | |
|---|---|
| Non-template strand | 5'-AGGCTTGACACTTTATGCTTCGGC(T/Cy3B)CGTATA ATGTGTGGAATTGTGAGAGCGGATAACAATTTC-3' |
| Template strand | 5'- GAAAT(T/Atto647N)GTTATCCGCTCTCACAATTCCA CACATTATACGAGCCGAAGCATAAAGTGTCAAGCCT-3' |
| **rrnB P1** construct for measurements in Figs. 4B, 4C, 5D-F | |
| Non-template strand | 5'-(biotin)CTCTTGTCAGGCCGGAATAACTCCCTATAAT GCGCCACCACTGACACGGAACAACGGCAAACAC-3' |
| Template strand | 5'- GTGTTTGCCGTTGTTCCGTGTCAGTGGTGGCGCATT ATA GGGAGTTATTCCGGCCTGACAAGAG-3' |
| **-15/+20 rrnB P1** construct for measurements in Fig. S13 | |
| Non-template strand | 5'-CTCTTGTCAGGCCGGAATAACTCC(T/Cy3B)TATAAT GCGCCACCACTGACACGGAACAACGGCAAACAC-3' |
| Template strand | 5'- GTGTTTGCCGTTGTTCCGTGTCAGTGGTGGCGCATT ATA AGGAGTTATTCCGGCCTGACAAGAG-3' |
| **pR** construct for measurements in Figs. 4A, 5A-C | |
| Non-template strand | 5'-(biotin)ATCTATCACCGCAAGGGATAAATATCTAACA CCGTGCGTGTTGACTATTTTACCTCTGGCGGTGATAAT GGTTGCATGTAGTAAGGAGGTGGTATGGAAT-3' |
| Template strand | 5'- ATTCCATACCACCTCCTTACTACATGCAACCATTAT CACCGCCAGAGGTAAAATAGTCAACACGCACGGTGTT AGATATTTATCCCTTGCGGTGATAGAT-3' |

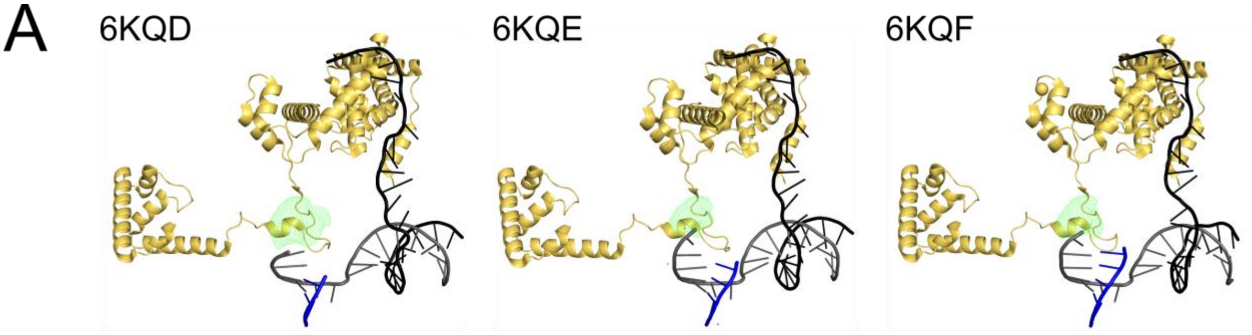


B

| Structure (PDB file) | $\langle R_{DA} \rangle_E$ | Fret efficiency, E ($R_0$=60Å) |
|---|---|---|
| RPo (4G7H) | 60.8±3.2 | 0.480 |
| RPitc3 (6KQD) | 61.0±3.1 | 0.474 |
| RPitc4 (6KQE) | 60.6±3.0 | 0.485 |
| RPitc5 (6KQF) | 61.1±3.0 | 0.473 |
| RPitc6 (6KQG) | 60.8±2.4 | 0.480 |
| RPitc7 (6KQH) | 60.7±2.5 | 0.482 |

**Fig. S1.** Accessible volume measurements for labelling the base of the σ-finger.

(A) Accessible volume modelling of fluorescent probe Cy3B placed at position $\sigma^{70}$ residue 511 ($\sigma^{A}$ residue 321) on complexes with 3 (6KQD), 4 (6KQE) and 5 (6KQF) nucleotides of RNA. $\sigma^{70}$ is straw coloured; accessible volume modelling shown is green; RNA is in blue; template DNA in gray, and non-template DNA in black.

(B) Accessible volume distance measurements between a label at the base of the σ-finger ($\sigma^{A}$ residue 319) and $\sigma^{70}$ residue 366 ($\sigma^{A}$ residue 174) showing that a label base of the σ-finger is not sensitive to movements at the tip of the σ-finger. $\sigma^{A}$ residue 319 was used for these comparisons as $\sigma^{A}$ residue 321 was not present in PDB files 6KQG and 6KQH.

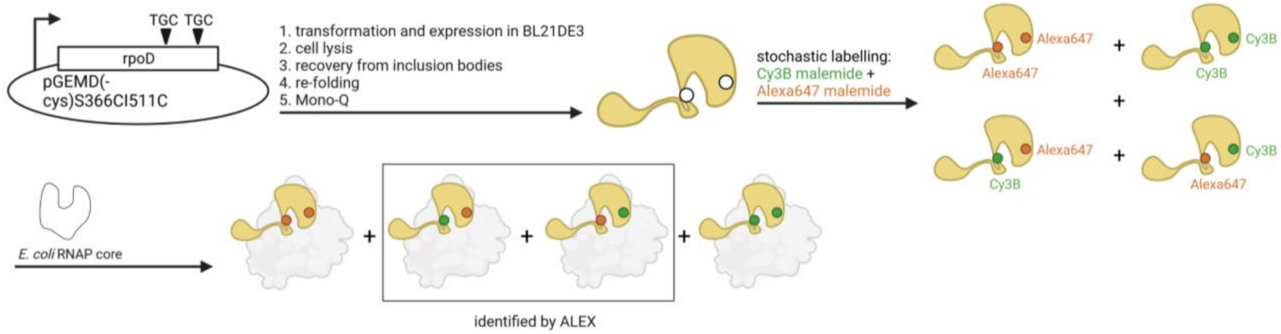


**Fig. S2.** Summary of the labelling method used to generate a DL RNAP-$\sigma^{70}$ construct labelled at positions 511 and 366 of $\sigma^{70}$ with dyes Cy3B and Alexa647.

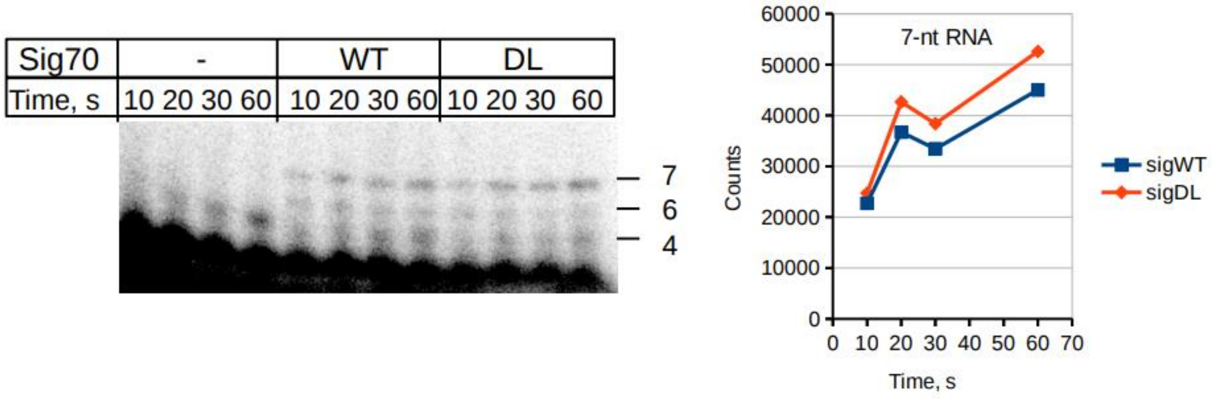


**Fig. S3.** Functional assays of double labelled $\sigma^{70}$ derivative. *In vitro* transcription assay comparing initial transcription profiles obtained using the DL RNAP-$\sigma^{70}$ and a wild type RNAP-$\sigma^{70}$.

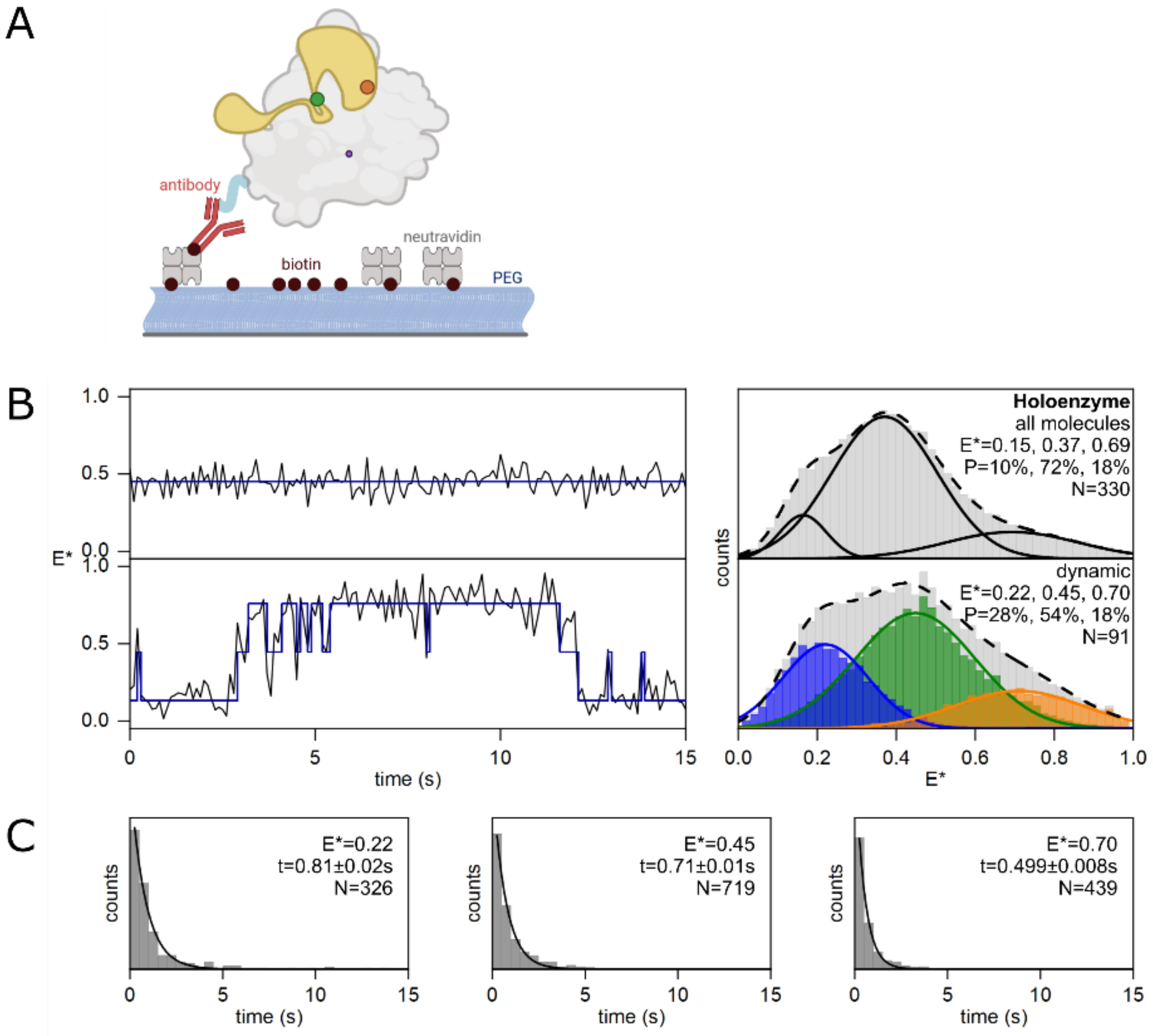


**Fig. S4.** *In vitro* smFRET results with DL RNAP-$\sigma^{70}$.

(A) DL RNAP-$\sigma^{70}$ was immobilized via a hexahistidine tag on a glass slide functionalized with anti-hexahistidine tag antibody.

(B) smFRET data for the σ-finger in DL RNAP Holoenzyme complexes showing static (*upper*) and dynamic (*lower*) behaviour. Left, representative traces of static and dynamic behaviour. Right, E* histograms formed as a result of hidden Markov modelling, and Gaussian fitting of sub-populations.

(C) Dwell time histograms of each of the states found by hidden Markov modelling of traces exhibiting dynamic behaviour.

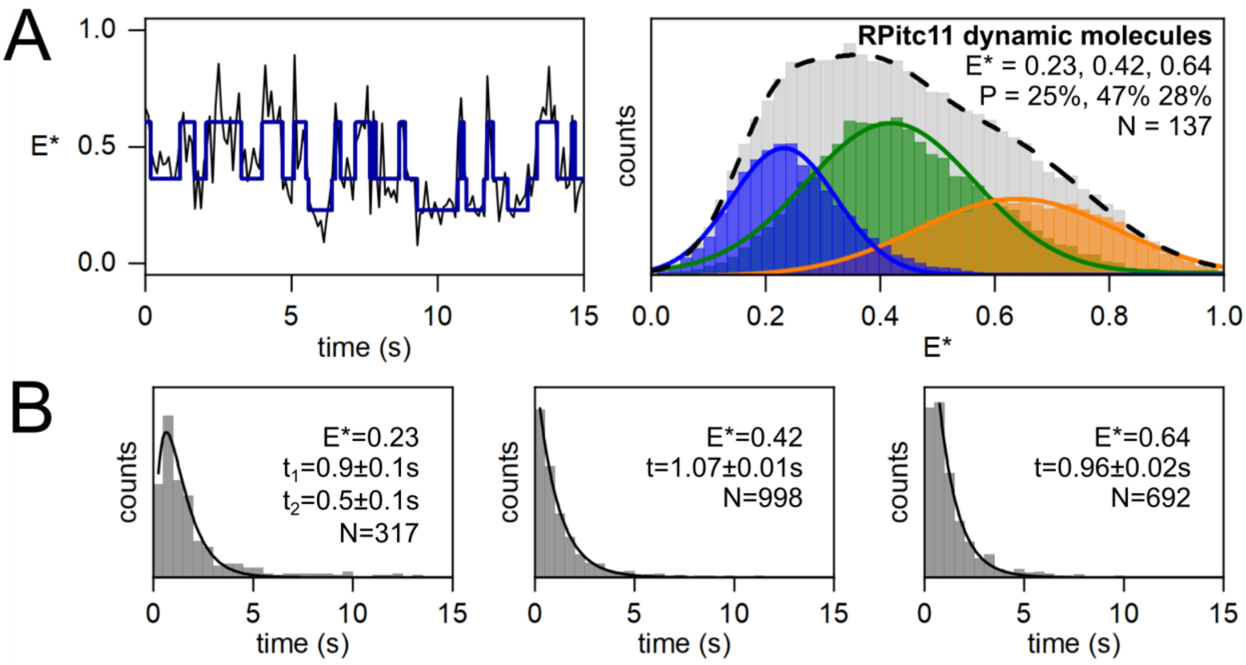


**Fig. S5.** *In vitro* smFRET data for dynamic σ-finger $RP_{itc11}$ molecules formed with lacCONS promoter and ApA initiating dinucleotide.

(A) Left, representative traces of dynamic behaviour. Right, E* histograms formed as a result of hidden Markov modelling, and Gaussian fitting of sub-populations.

(B) Dwell time histograms of each of the states found by hidden Markov modelling of traces exhibiting dynamic behaviour.

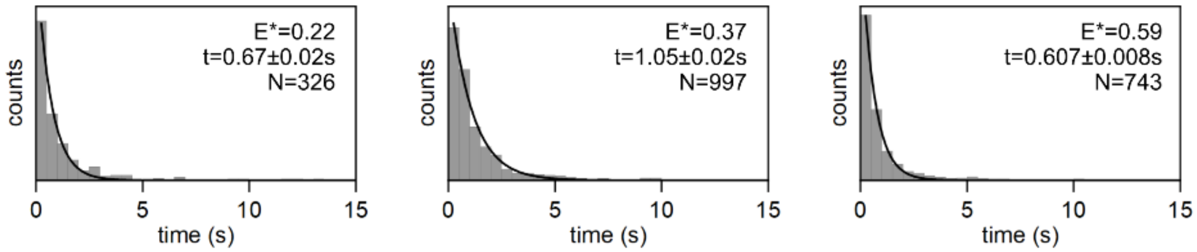


**Fig. S6.** Dwell time histograms for $RP_{itc2}$ molecules formed with the lacCONS promoter and ApA initiating dinucleotide. States are found by hidden Markov modeling of individual smFRET trajectories exhibit conformational dynamics of the σ-finger.

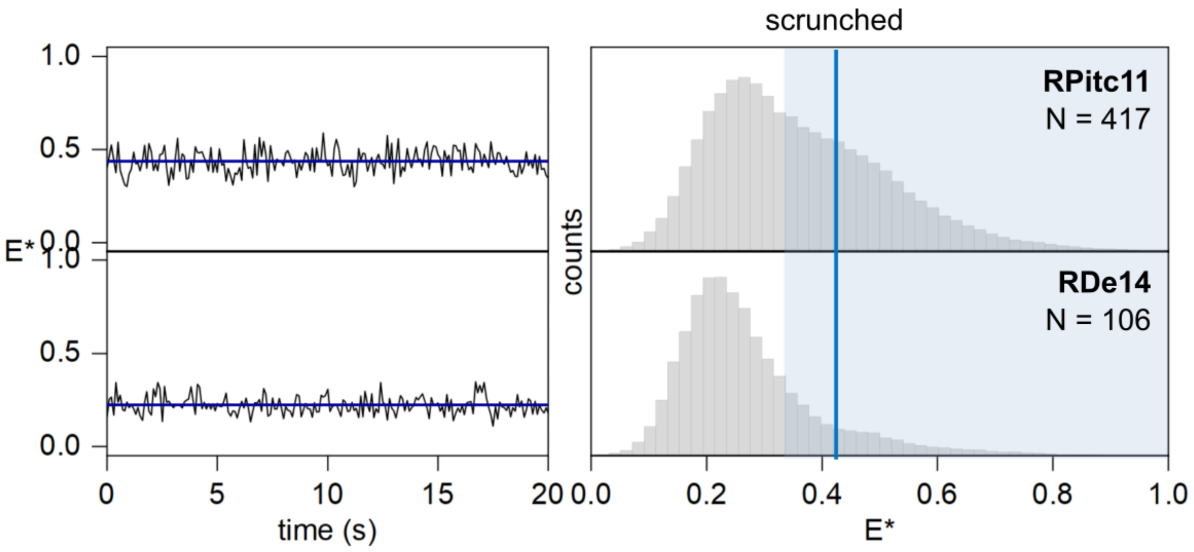


**Fig. S7.** smFRET data monitoring the conformation of the DNA transcription bubble in $RP_{itc11}$ (*upper*) and $RD_{e14}$ (*lower*) complexes formed with lacCONS promoter labelled at positions -15 (non-template DNA) and +20 (template DNA) and ApA initiating dinucleotide. Left, representative traces. Right, E* histograms.

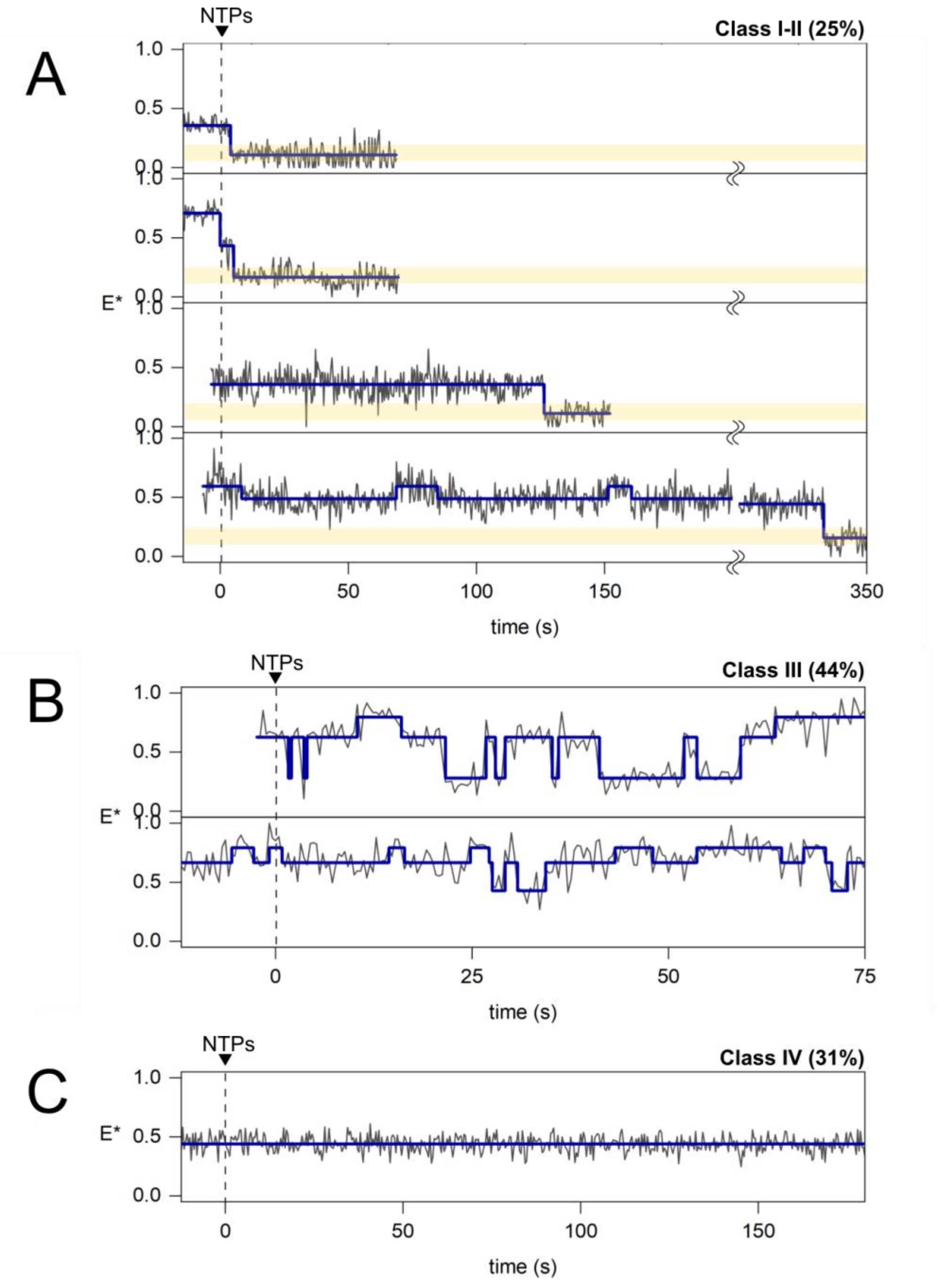


**Fig. S8.** Example traces of each class of E* time trajectories obtained in real-time σ-finger experiments involving the lacCONS promoter and ApA initiating dinucleotide. See text for detailed description of the 4 classes.

(A) Class I and Class II. The conformation after displacement is highlighted in yellow.

(B) Class III.

(C) Class IV.

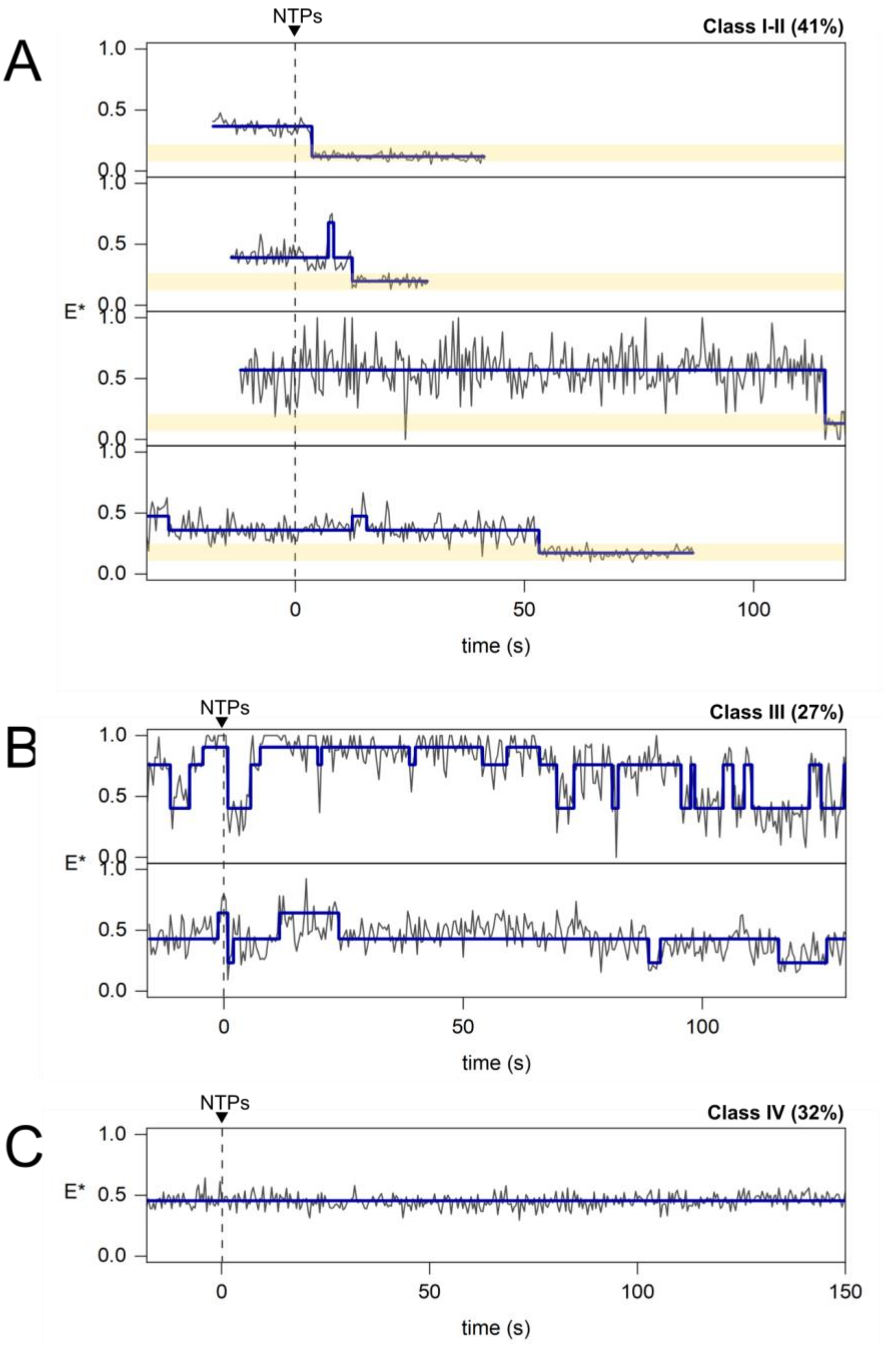


**Fig. S9.** Example traces of each class of E*-time trajectories obtained in real-time σ-finger experiments involving the lacCONS promoter and pppApA initiating dinucleotide.

(A) Class I and Class II. The conformation after displacement is highlighted in yellow.

(B) Class III.

(C) Class IV.

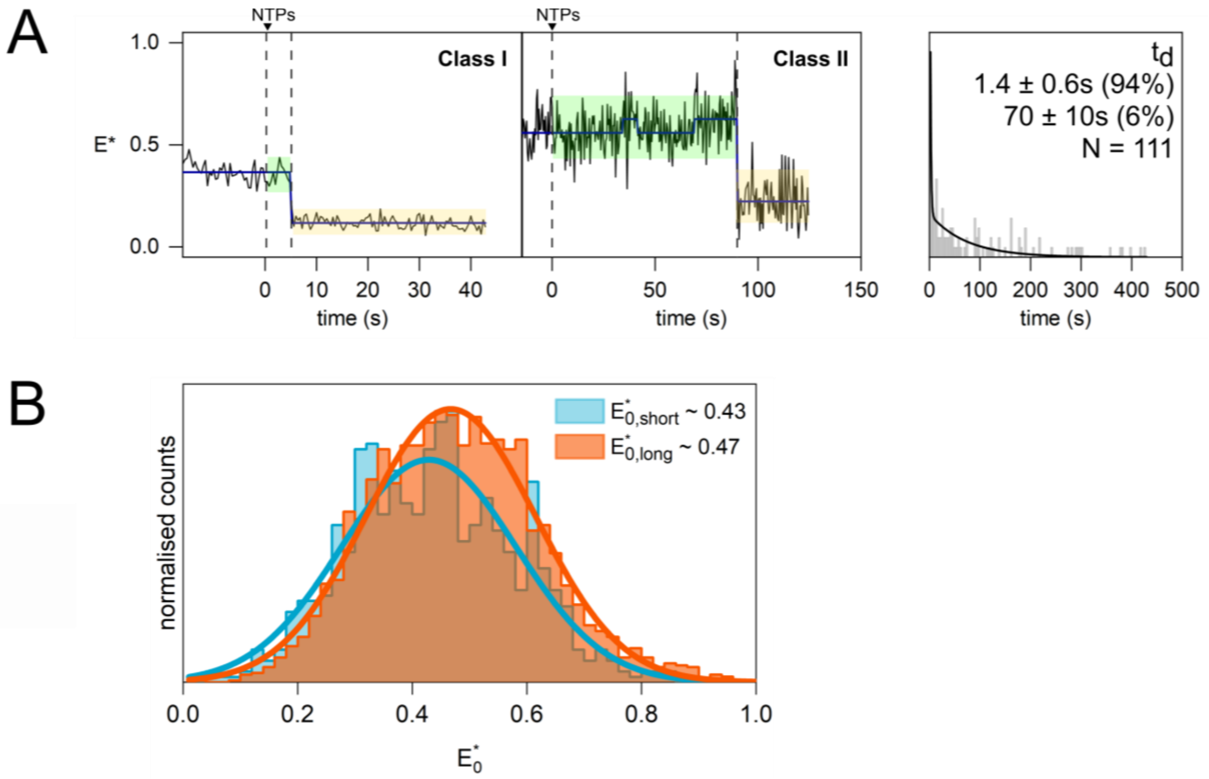


**Fig. S10.** smFRET data tracking the σ-finger for the lacCONS promoter and synthesis of RNA with a 5'-triphosphate end.

(A) Representative E*-time trajectories and dwell time histogram for the time to displacement, $t_d$, fitted to a double-exponential decay (black line). The conformation between NTP addition and displacement is highlighted in green, and the conformation after displacement is highlighted in yellow.

(B) Histograms showing the σ-finger conformation before NTP addition of Class-I ($E^*_{0,short}$; blue) and Class-II ($E^*_{0,long}$; red) molecules.

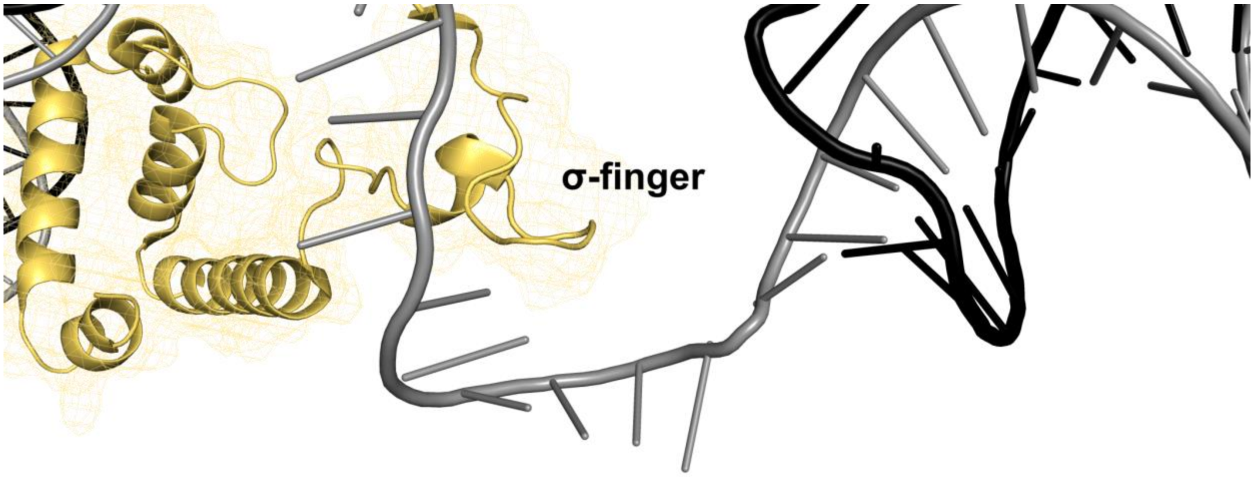


**Fig. S11.** Structure of the pR $RP_o$ structure (7MKD). The $\sigma^{70}$-factor is straw coloured; template DNA is in grey, and non-template DNA in black. The σ-finger is estimated to clash with a growing RNA chain of 4- to 6-nt in length.

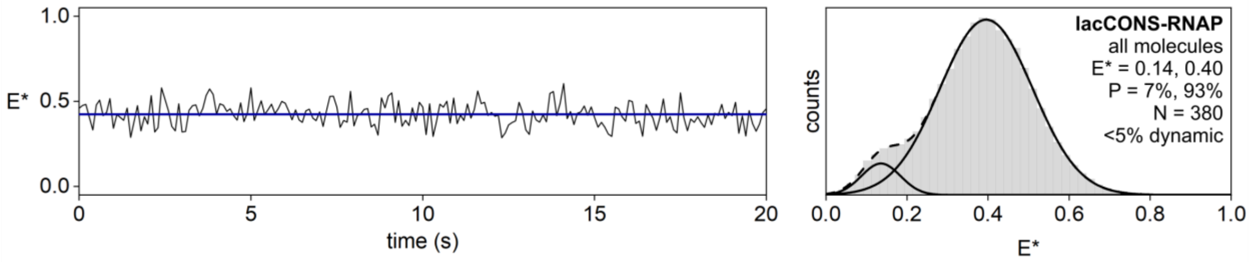


**Fig. S12:** smFRET data for the lacCONS DL RNAP-$\sigma^{70}$ complex. Left, representative trace. Right, E* histogram with bi-modal Gaussian fitting (black line).

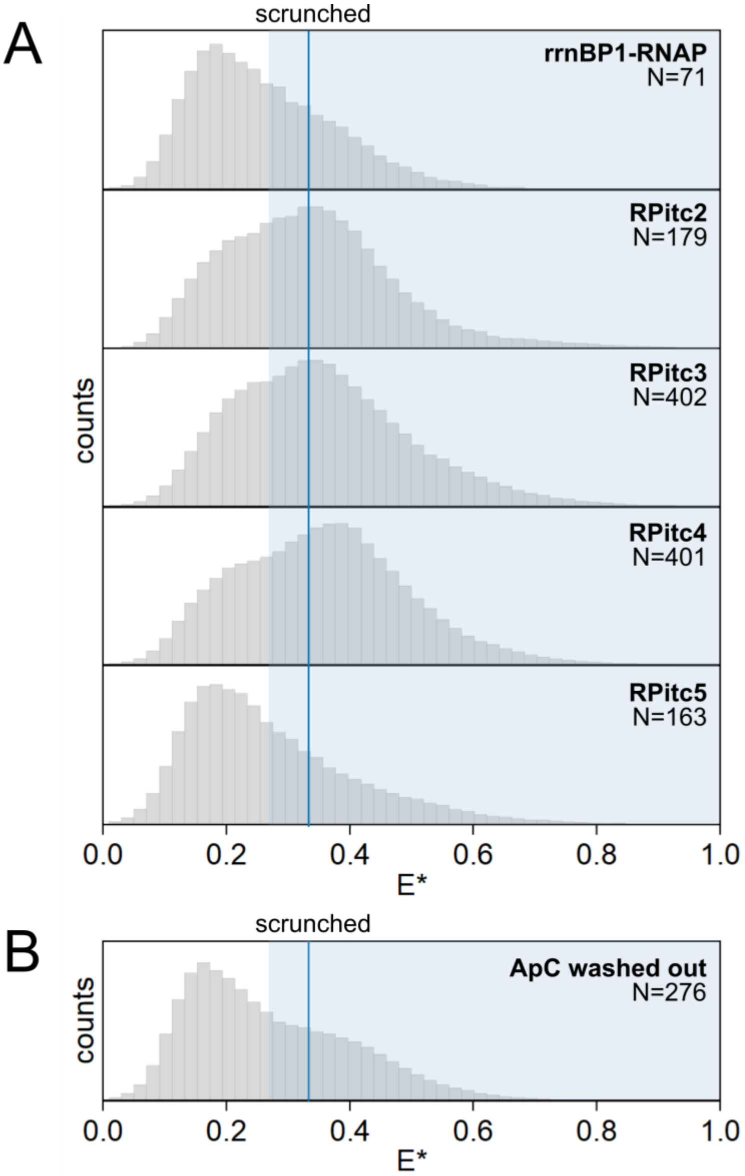


**Fig. S13**. smFRET data monitoring the conformation of the DNA transcription bubble in complexes formed with the rrnBP1 promoter labelled at -15 (non-template DNA) and +20 (template DNA) and ApC initiating dinucleotide.

(A) E* histograms of the rrnBP1-RNAP complex, $RP_{itc\leq2}$, $RP_o$, $RP_{itc\leq3}$, $RP_{itc\leq4}$ and $RP_{itc\leq5}$

(B) E* histogram of the complex formed after forming $RP_{itc\leq2}$ and subsequent washing out of ApC.
